# Yes1-mediated Cul9 phosphorylation promotes the metabolic reprogramming in gastric cancer

**DOI:** 10.1101/2023.10.18.562906

**Authors:** Youliang Wu, Heng Zhang, Shangxin Zhang, Mingliang Wang, Huizhen Wang, He Huang, Xuehui Hong, Zhiyong Zhang, Yongxiang Li

**Affiliations:** Department of General Surgery, the First Affiliated Hospital of Anhui Medical University, Hefei 230022, People’s Republic of China; Department of Histology and Embryology, Xiang Ya School of Medicine, Central South University, Changsha, Hunan, China; NHC Key Laboratory of Carcinogenesis, Cancer Research Institute and School of Basic Medicine, Central South University, Changsha, Hunan, China; Department of Gastrointestinal Surgery, Zhongshan Hospital of Xiamen University, Xiamen, Fujian, China; Department of Surgery, Rutgers University Hospital, Robert-Wood-Johnson Medical School, New Jersey, USA

**Keywords:** Cul9, Yes1, gastric cancer, *Helicobacter pylori*, nucleotide metabolism

## Abstract

Although Cul9 has been implicated in human carcinogenesis, its upstream regulators and roles remain unknown. Herein, we indicate that the Cul9 promoter is hypermethylated in GCs. Bioinformatics, mass spectrometry, and unbiased-kinase screen identify the tyrosine kinase Yes1 as a key regulator of Cul9. Yes1 phosphorylates Cul9 at Y1505, promoting its selective autophagy. Patient-associated mutation of Yes1 or helicobacter pylori infection induces Cul9-Y1505 phosphorylation which switches Cul9 from a tumor-suppressor to an oncogene, as evidenced by the fact that Cul9-Y1505D knockin mice are more susceptible to gastric tumorigenesis than wild-type counterparts. Metabolic profiling and ATAC sequencing reveal that Cul9-Y1505D mutant promotes pyrimidine and purine synthesis pathways in GC. DNA-demethylating drug decitabine or HG78 compound upregulates Cul9 expression and limits GC cell proliferation in a Yes1-dependent manner. The Yes1 inhibitor CH6953755 or Leflunomide and Mycophenolate mofetil (MMF) also impair the malignancy of GC with Cul9 dysregulation. Cul9 in turn binds Yes1 and disrupts Yes1 stability, establishing a feedback circuit. Collectively, our project reveals an unrecognized role of the Yes1-Cul9 loop in GC, suggesting potential therapeutic targets.

## Introduction

Cullins (*cul1*, *2*, *3*, *4A*, *4B*, *5, 7, and 9*) constitute a distinct E3 ubiquitin ligase family by binding to ROC1/RBX1, a small RING finger protein [1]. Relative to other family members, Cul9, also named the p53-associated, PARkin-like cytoplasmic protein (PARC), shares extensive sequence homology with Cul7 [2]. It is well-known that the Culs family took part in carcinogenesis [3]. But the role and mechanism of Cul9 in tumor progression are not clear so far. For example, in acute myeloid leukemia (AML), cul9 was previously found to block the nuclear translocation of tumor suppressor p53 and antagonized p53-induced apoptosis [4], suggesting its oncogenic role. However, this finding does not seem to be reproducible based on the findings: 1), In *Cul9 knockout* mice, the functions and localization of p53 did not change [5]; 2), in most cells with or without Cul9 and p53 overexpression, Cul9 did not induce the cytoplasmic accumulation of p53 [6]. On the contrary, these studies indicate that cul9 acts as a novel activator of p53 and inhibits lymphomagenesis [6, 7]. Therefore, it is still a challenge to clarify the functions of Cul9 in leukemia. Recent data also suggest the effects of Cul9 in solid tumors. However, very little is known about this aspect.

Mutations or dysregulation of Src tyrosine kinases (SFKs) family takes part in oncogenesis [8]. The SFKs include eight members: Fyn, Src, Yes1, Hck, Lyn, Fgr, Lck, and Blk. Because the SFKs dictate intracellular signal transduction pathways in various diseases, they are excellently druggable targets [8]. Although Fyn, Src, Yes1, and Lyn are detected in almost all cell types, the expression of other members is tissue-specific [9]. Tyrosine residue autophosphorylation at Y419 in the activation loop enhances the kinase activity of SFKs, while the phosphorylation of Y529 residue at the C-terminus suppresses the catalytic ability of SFKs [10]. Beyond phosphorylation, ubiquitination also modulates the activity of SFKs. For example, c-Cbl E3 ligase degrades several SFKs (Src, Fyn, Yes1, Lyn) by ubiquitination, thereby inhibiting the ability of cell transformation induced by SFKs [11]. In turn, the activation of SFKs also affects the ubiquitination of c-Cbl itself and its substrates [12], indicating a reciprocal regulation between E3 ligase and SFKs. We previously reported the oncogenic function of Yes1 in gastric cancer (GC) [13]. However, the crosstalk between Yes1 and E3 ligase in GCs has not been defined.

Here, Yes1 is identified as a previously unknown substrate of Cul9 E3 ligase. We also demonstrate that Cul9 is a Yes1-dependent tumor suppressor in GC. High-throughput screen identifies HG78 compound as an activator of Cul9, which limits GC development by targeting MBD2, a CpG methylation reader. These novel findings suggest that targeting the Yes1-Cul9 loop represents a promising therapeutic strategy in treating GC.

## Materials and Methods

### Reagents, constructs, and cell culture

Protease inhibitor MG-132 (Cat#: 1211877-36-9) and cocktail (Cat#: sc-45045) were bought from Santa Cruz. Yes1 inhibitor CH6953755 was purchased from MedChemExpress. Cycloheximide (CHX), Sodium vanadate, Tween 20, and X-film were got from Sigma-Aldrich or Millipore, respectively. Leflunomide and Mycophenolate mofetil (MMF) were bought from AET PHARMA and Roche, respectively. The library containing 416 compounds for kinase screening was obtained from Selleck Chemicals (Houston). PBS, antibiotics, Lipofectamine 2000, HEPES, and Glutamine were bought from Invitrogen. Site-Directed Mutagenesis Kit (Catalog #210515 and #210513) was got from Agilent. Antibodies were purchased from the following sources: Cul9 (Bethyl, A300-096A), MBD2 (Abcam), LC3B (Abcam, ab192890), p62 (ab109012). Anti-GST antibody (Abcam #ab111947 and Sigma, #G7781-100UL), anti-His (Abcam, #ab18184), anti-MYC (A190– 205A), anti-COP1 (Abcam, ab56400), anti-Flag M2 (Sigma, F3165-2MG), Ki-67 (Cell Signaling, #9129), OPTN (Cell Signaling, #70928), anti-Yes1 (Abcam, ab109265), Tollip (Sigma, #HPA038261), NDP52 (Abcam, #ab68588), NIX (Abcam, #ab8399), NBR1 (Abcam, #ab55474), anti-HA (Sigma, #H6908-100UL), anti-Tubulin (A302-630A), and anti-GFP (Abcam, #ab290) were provided by Dr. Bing Wang. Anti-ub-K63 and β-actin antibodies were bought from Millipore. Secondary antibodies were got from ThermoFisher (Cat#G-21234) and Bio-Rad (#1706515 and #1706516). p-Cul9-Y1505 antibody was obtained by a similar method previously reported [14]. Subcloning of the *Cul9* promoter including the MBD2 methylation target sequence was performed using a pCpGI luciferase reporter without CpG (Life Technologies). *Cul9*, *COP1*, and *Yes1* plasmids were from Addgene. Efficient methods for constructing *MBD2*, *Cul9-pCPGI* without CpG luciferase reporter methylation promoters, and mtMBD2 plasmid without methylation domains) have been indicated. Normal gastric mucosal epithelial cells (GES-1), AGS, MKN45, SGC7901, BGC823, HEK293, and HGC27 cell lines were from ATCC.

### Chromatin immunoprecipitation (ChIP)

ChIP assays were carried out with the ChIP kit (Millipore, USA) and anti-MBD2. Polymerase chain reaction (PCR) was used to amplify immunoprecipitated DNA using the Cul9 promoter combined with the indicator primer of the CpG islands. Transcriptional activation activity of Cul9 was detected by applying a luciferase kit (Promega).

### Methylated CpG-DNA immunoprecipitation

In the previous description [15], the process of immunoprecipitation methylated CpG-DNA was performed. And with an ABI StepOne plus PCR instrument, the methylated DNA was assayed by using sheared DNA and PCR analysis.

### Luminescence high-throughput screen

HEK293 cells carrying an inactive MDB2-luciferase reporter were used for a luminescence-based high-throughput screen. All screened compounds were bought from Cayman, Chembridge, and Selleckchem chemicals, respectively. Individual compounds from the chemical library that are suitable for dissolution in DMSO will be dissolved in advance. Drug plates for screening were prepared with a medium at a final concentration of 1 μM. One day before compound treatment (1 μM of compounds/well, n=3), approximately 20,000 cells were seeded into each well. 3 days after small molecule treatment, cells were washed with PBS, and then co-cultured with Bright-Glo (Promega). Fifteen minutes later, the luminescence of activated MBD2-luciferase was measured using a MicroBeta Trilux (Perkin Elmer) microplate reader. An active compound was considered by a two-fold or more change in the relative luciferase unit.

### Knockdown assays

Knocking down *Cul9*, *COP1*, *Yes1*, *ATG5*, *p62*, *BECN1*, *MBD2*, and *OPTN* was performed according to the protocol of Invitrogen, Santa Cruz Biotechnologies, and Sigma, respectively. Transfection was performed with Lipofectamine 2000 or FuGENE® 6. Real time-PCR and Western blots were used to verify these transfections. Their sequence was as follows: *siCul9-*#1: 5′-GCUGAGAGACACGUUGUUUAG-3′; *siCul9-*#2: 5′-UACUGAGGGUGCUCUUCUG-3′; or *scramble*: 5′-GAUGCUCGCACAGCACAAU-3′; *shYes1-#1*: 5′-CCAGCCTACATTCACTTCTAA-3′ and shYes1-#2: 5′-CCTCGAGAATCTTTGCGACTA-3′; *shcontrol*: 5′-CCGCAGGTATGCACGCGT-3′. *shp62*: 5’-TAGTACAACTGCTAGTTATTT-3’; the targeting sequence of *BECN1* is 5’-CCGACTTGTTCCTTACGGAAA-3’, and that of *OPTN* is< 5′-GCACGGCATCGTCTAAATA-3′. siRNA against MBD2: 5’-GAAGAUGAUGCCAGUAAUUUU-3′ and its control: 5′-UUCUCCGAACGUGUCACGUTT-3′. *siCOP1* and shRNAs against *Cul9* and *ATG5* were from Origene, shCOP1 was from Santa Cruz Biotechnology, and siRNA against Yes1 was from Invitrogen.

### TCGA data analysis and ATAC sequencing

Whole genome RNA sequencing (RNA-seq) data on the TCGA dataset for gastric cancer can be downloaded from the following website: https://xenabrowser.net. Patients for whom survival data were not available were excluded. Relevant clinical characteristics were obtained from the TCGA dataset, including gender, age, pathological TNM, survival time, etc. According to the Cul9 gene expression level, the samples were divided into three groups: low, medium, and high. The cutoff points are represented by the 25th and 75th percentiles. The false discovery rate was calculated by Benjamini–Hochberg procedure, and the top 500-upregulated genes for gene ontology analysis was described before.

### EdU assay

The effect of Cul9 on GC cell proliferation was assessed by EdU assay using an In Vivo EdU Click kit (Sigma) according to the manufacturer’s instructions. Briefly, GC cells treated with 10 μM EdU were cultured for 12 h. Then, cells were fixed with 4% paraformaldehyde and incubated with Triton X-100 and BSA (0.3%) in PBS. Cells were stained with the Click Additive Solution provided in the kit and DAPI, respectively. A fluorescence microscope (Nikon) was used for image capturing, and ImageJ software was used for fluorescence signal analysis. Finally, the percentages of EdU+ cells were analyzed.

### Proximity Ligation Assay (PLA)

Briefly, GC cells were fixed with paraformaldehyde (4%), and Triton X-100 (0.1%) was for permeabilization. Blocked about 30 min later in the PBS buffer with 5% BSA and 5% donkey serum and 0.3% Triton X-100, the manufactory’s instruction was followed with Duolink® In Situ Detection Reagents (Sigma-Aldrich). The interaction between proteins was found using Image J software.

### GC samples and Immunohistochemistry (IHC)

All histologically confirmed GC specimens and adjacent normal gastric specimens were gathered from the general surgery department of the first affiliated hospital, Anhui medical university between 2006 and 2008. These samples were used to construct tissue microarray (TMA, Shanghai Outdo Biotech Company). All participating patients were given informed consent and had complete clinical follow-up data. The ethics of this study was supported by the Review Ethics Committee of Anhui Medical University. TMA was used to analyze CUL9 and other related protein expressions by the immunohistochemical (IHC) method. Detailed IHC steps and immunoreactivity scoring (IRS) methods were described in previous studies [16, 17]. Briefly, IHC was performed on TMA to analyze the staining intensity and percentage of mucosal cells. The result of multiplying these two factors was defined as IRS. Patients were defined by IRS as low expression group (-, IRS<4) and high expression group (+, IRS ≥4) by IRS.

### Cell incubation and target expression construction

All the cell lines we used in the study were were incubated in RPMI-1640 (Gibco) with 10%FBS (Gibco) at the humidified condition of 37°C with 5% CO2. To get transient and stable indicated cells, appropriate cells were transfected or infected with different target gene expression plasmids or lentiviral (as shown in figures), respectively, following the manufacturer’s protocol.

### Immunoprecipitation (IP) and immunoblotting (IB)

IP and IB assay methods were shown previously to evaluate protein interaction [18, 19]. To obtain enough cell samples, we inoculate the indicated cells in a 10cm petri dish and collect the cell samples when the confluent degree reaches 80%. Briefly, before harvest, the indicated samples were lysed on ice using the indicated buffer (including a cocktail of phosphatase and protease inhibitor) for 30 minutes. Subsequently, crude suspensions lysate was removed by centrifugation at 12000 g for 10 min at 4°C, and then the supernatant was treated with appropriate beads. After that, the beads were cleaned with lysis buffer to eliminate unbound protein, and the immune complexes bound to the beads were boiled for eight minutes for elution, followed by IB analysis. Cell lysates and immune complexes were loaded onto 6–15% SDS-PAGE (based on the molecular weight of the protein shown) for separation and then imprinted on the PVDF membrane to incubate the appropriate primary antibody (as shown in figures) and corresponding secondary antibody. Finally, the protein signals on the membrane were quantified with an ECL detection kit.

### Quantitative real-time PCR (qRT-PCR)

Total RNA from differently treated cells was extracted with Trizol Reagent (Invitrogen). Subsequently, the extracted total RNA was reverse transcribed into cDNA and amplified on an Applied Biosystems in triplicate in SYBR-Green Mix (Toyobo), respectively. β-actin gene was regarded as normalized, and all target gene primers were listed as follows: β-actin, 5ʹ-TAACCAACTGGGACGATATG-3ʹ (F) and 5ʹ-AAACAGGGA CAGCACAGCCT-3ʹ (R); Cul9, 5′-AACCCTGGAACAGAAGAG-3′ (F) and 5′-GAGAGGACATCTGTACTGG-3′ (R); MBD2: 5ʹ-AGCAAGCCTCAGTTGGCAAGGT-3′ (F) and 5′-TGTTCATTCATTCGTTGTGGGTCTG-3′ (R); GAPDH: 5ʹ-ACTCCACTCACGGCAAATTC-3′ (F) and 5′-TCTCCATGGTGGTGAAGACA-3′ (R); Yes1: 5ʹ-GCTAGAACTACAGAAGACCTTTCA-3′ (F) and 5′-TCTCGAGGGATTTCCCAAGCATCT-3′ (R); COP1: 5ʹ-GAGCACCGGATCTGGATAAA-3′ (F) and 5′-ACATGCCGAGCTGAAGTTTT-3′ (R) ; p62: 5′-TCCTGCAGACCAAGAACTATGACATCG-3′ (F) and 5′-TCTACGCAAGCTTAACACAACTATGAGACA-3′ (R); OPTN: 5′-GCCTCCGCGGATTCGAAATGTCCCATCAACCTCTCAGCT-3′ (F) and 5′-TTCAATCGATGTTCGAATTAAATGATGCAATCCATCACGTGAATCTG-3′ (R); ATG5: 5′-TGTGCTTCGAGATGTGTGGTT-3′ (F) and 5′-GTCAAATAGCTGACTCTTGGCAA-3′ (R); DHODH: 5′-AGAGAGCTGGGCATCGAC-3′ (F) and 5′-AACCCCGATGATGGGAAT-3′ (R); ATIC: (F) 5′-TTGGAGACTAGACGCCAGTTA-3′ and 5′-GGCATCTGAGATACGCCTTTG-3′ (R); BECN1: 5′-ATGGAGGGGTCTAAGGCGTC-3′ (F) and 5′-TCCTCTCCTGAGTTAGCCTCT-3’ (R).

### Cell proliferation, migration, and invasion assays

Cell growth and proliferation capabilities of different indicated cells were assessed by cell proliferation and colony formation assay. Briefly, for the cell proliferation assay, differently indicated Cell viability was determined with CCK8 at appropriate time points. As for colony formation assay, indicated cells were seeded in 6-well plates for about two weeks to form visible colonies. After crystal violet staining, the results were collected. For migration assay, different indicated cells in 200 μl serum-free medium were inoculated to the upper Transwell chamber with 8-μm pores, and then 600 μl medium with 10% FBS was loaded into the lower chamber for chemotactic. Upper chamber (non-migrated) cells were removed and low chamber (migrated) cells were fixed and stained with 0.5% crystal violet after 24 hours, and migrated cells number was counted under a light microscope. In addition, cell invasion assay was evaluated in matrigel-coated transwell chambers. Use a similar method to the migration assay described above.

### Immunofluorescence (IF)

Briefly, the indicated cells were plated in 6-well plates with pre-placed glass coverslips. Firstly, when the confluence of cells reached 70% under appropriate treatment conditions, they were fixed, permeabilized, and blocked with 4% PFA, 0.2% Triton X-100 and 1% BSA in PBS, respectively. Then the cells were incubated with appropriate antibodies and stained with/without DAPI as needed. Finally, the cell images were scanned and analyzed using a confocal microscope (Carl Zeiss, Germany).

### Xenograft model and Knockin mice

The animal experiment in this study was carried out in compliance with NIH guidelines and approved by the Animal Care and Use Committee of Anhui Medical University. The xenograft model in vivo was used to further study the Cul9 protein function. For subcutaneous xenograft models, immune-deficient BALB/c nude mice (5-week-old) were injected with indicated cells in the right or left flank subcutaneously. At an appropriate time, nude mice were sacrificed and the tumor body was taken out for evaluation of tumor volume [V = (depth × width × length)/2], mass, and indicated protein expression. Furthermore, we constructed WT-Cul9 and Cul9-Y1505D knockin gp130F/F mice to further explore the role of Cul9 in the development and progression of gastric cancer in vivo. When appropriate, these mice were collected and analyzed for stomach size, tumor mass, tumor incidence, and survival time.

### In vitro ubiquitination assay and mass spectrometry (MS) analysis

In brief, for in vitro ubiquitination assays, endogenous Cul9 was expressed in AGS cells and purified with anti-Cul9 antibody; Yes1 was purified from AGS cell lysates with anti-Yes1 antibody after transfected with different indicated recombinant Yes1 plasmids. Then, purified substrate Cul9 and different indicated Yes1 were incubation in a ubiquitination reaction buffer (1 mM DTT, 50 mM Tris–HCl, 5 mM MgCl2, 30μM MG132, pH 7.5, 1 μM HA-Ub, 2mM NaF and 2 mM ATP). When the reaction was completed, the mixture reaction buffer was IP with anti-Yes1 beads, and the immunoblotting with different indicated antibodies was used to analyze the Yes1 protein ubiquitination level. For MS analysis, the above samples were separated on 10% SDS-PAGE gel and stained with silver. Target Yes1 ubiquitination bands were clipped and subjected to LC-MS/MS sequencing analysis to determine Yes1 ubiquitination sites.

### In vivo ubiquitination assay

Target cells were treated using different indicated constructs, including control siRNA or siCul9, Flag–Yes1, and HA-ubiquitin, and then treated with or without MG132 (5 μg/ml, Selleckchem, S2619) for 6h. After the appropriate treatment, the indicated cells were lysed with denaturing lysis buffer (0.1 M Na2HPO4 ∕ NaH2PO4, 10 mM b-mercaptoethanol, 0.01 M Tris–HCl, pH 8.0, 6 M guanidinium-HCl, 5 mM imidazole) and then centrifuged to obtain supernatant. The supernatant was purified by incubation with pretreatment of A&G-conjugated Flag target antibody beads. Subsequently, the immune complex bound to the beads was rinsed with a washing buffer and eluted with a boiling SDS loading buffer. Finally, eluted proteins were examined by the immunoblotting analysis with targeted antibodies.

### In vitro kinase assay

One μg active or inactive recombinant Yes1 protein was incubated at 30°C with purified recombinant WT-GST-Cul9 or Y1505-GST-Cul9 mutant protein in a 1x kinase reaction buffer (10 μM ATP and 0.2 mM Na3VO4) for 30 min. Then the kinase reaction assay was terminated by SDS sample buffer. Mass spectrometry and immunoblotting assay were used to analyze phosphorylation sites and phosphorylation levels, as previously described.

### Cycloheximide-chase assay

Cycloheximide (CHX) is a protein biosynthesis inhibitor used to analyze the half-life of the target protein. Briefly, for cycloheximide-chase assay, indicated samples were incubated with cycloheximide at a concentration of 100 μg/mL (C7698, Sigma-Aldrich) and the protein samples were obtained with a lysis buffer at appropriate time points. Finally, the protein samples were analyzed by immunoblotting with a targeted antibody.

### GST affinity-isolation assay

GST affinity-isolation assay is used to study the direct interaction between protein-protein. Briefly, for GST affinity-isolation assay, GST and GST–tagged fusion protein were induced by IPTG during Escherichia coli production, and then incubated and fixed on glutathione-Sepharose beads for purification as bait proteins. Bait proteins were incubated with targeted prey proteins in a reaction buffer at 4°C for 2 h. Then the beads were rinsed with wash buffer and boiled for elution in SDS loading buffer, and then analyzed by immunoblotting and Coomassie staining.

### *H. pylori* growth and co-culture with GC cell lines

All wild-type (WT) Cag+ *H. pylori* strains (7.13, J166, and PMSS1) used in our study were performed on trypticase soy agar plates supplemented with 5% sheep blood for in vitro passage, as described in previous studies [20–22]. These bacterial strains grew for 18 hours under cultured in Brucella broth (Invitrogen) with 10% FBS (Sigma) at a condition of 37°C with 5% CO2. For in vitro studies, GC cell lines were co-cultured with *H. pylori* at a MOI of 50:1.

### Metabolites analysis

Intracellular metabolites were performed by LC-MS/MS method as described in previous studies [23, 24].

### Statistics

The statistical differences in quantitative data between different groups were evaluated by SPSS 17.0 or GraphPad Prism 8 software, and tested by Student’s t-test or one-way ANOVA method. Spearman’s correlation analysis method was used to analyze the correlation between different factors. Kaplan-Meier method combined with log-rank test was applied to survival analysis. *P* < 0.05 means the difference is statistically significant, in addition, *, ** and *** indicate *P* < 0.05, *P* < 0.01, *P* < 0.001, respectively.

## Results

### Methylation-mediated downregulation of *Cul9* level is tightly associated with an unfavorable prognosis in GC patients

TCGA data indicated that GC tissues expressed a lower level of *Cul9* than non-tumor specimens (Figure 1A-B). Clinically, a low level of Cul9 suggested a poor prognosis in GC patients (Figure 1C). Consistently, the normal GES-1 cells expressed higher Cul9 than all GC cell lines (Figure 1D).

**Figure 1.**
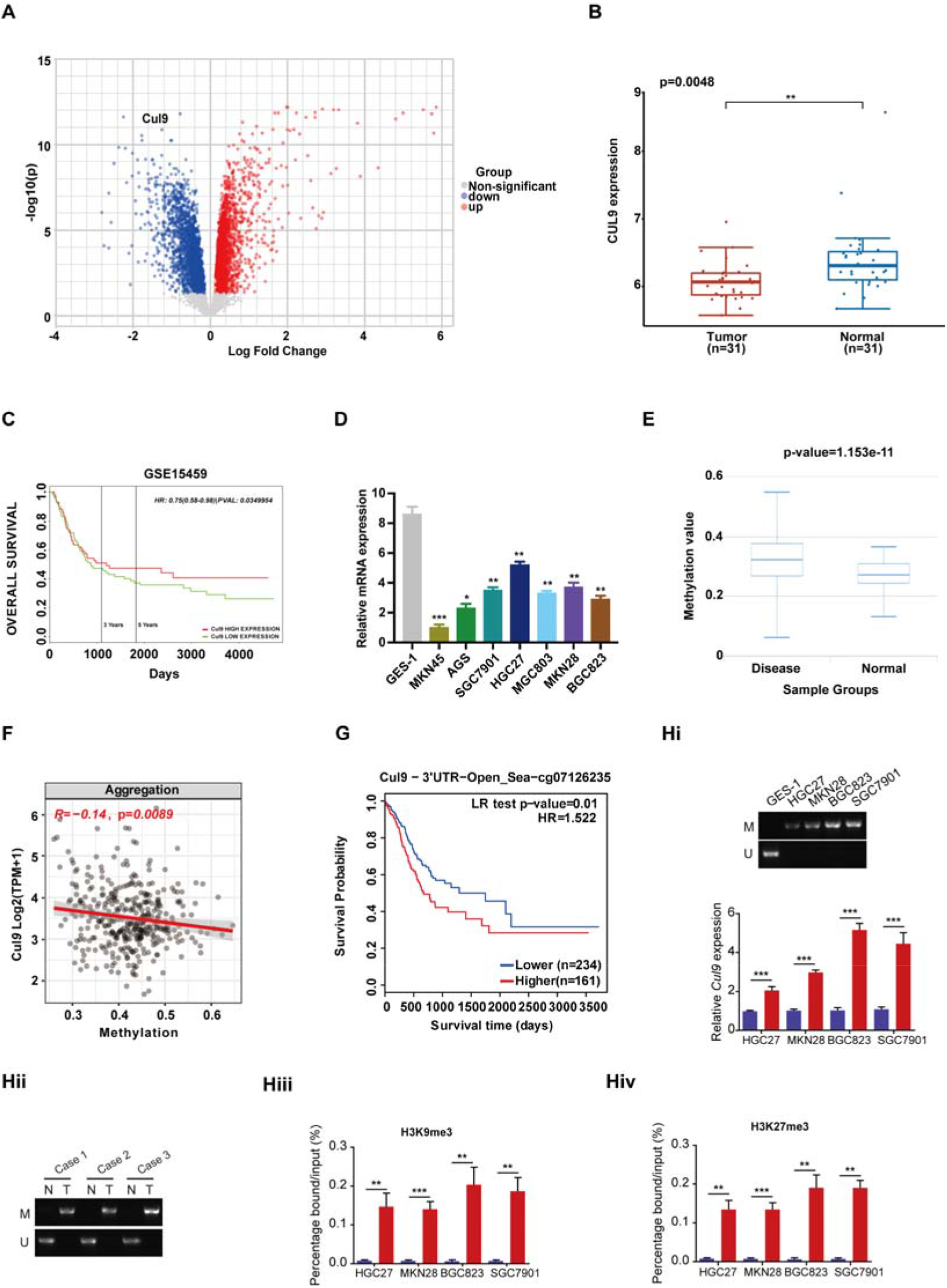
Methylation-mediated downregulation of the *Cul9* level suggests a bad outcome in GC patients. (A) Volcano plot of differential expressed genes from GC TCGA datasets. (B) Expression profiling of Cul9 mRNA in gastric cancers from GSE13911 datasets. Data shown are the log values of tumors (n=31) vs normal (n=31). (C) GC patients’ Kaplan-Meier assays based on the expression of Cul9. (D) The levels of *Cul9* were examined in indicated cell lines. (E) The CpG methylation in *Cul9* promoter using GC TCGA datasets. (F) The correlation between Cul9 levels and CpG. (G) The CpG methylation of *Cul9* and GC patients’ prognosis. (Hi) RT-PCR assays with normal and GC cell lines revealed the methylation of the *Cul9* promoter. (Hii) RT-PCR assays with GC patients and controls revealed the methylation of the *Cul9* promoter. (Hiii-iv) ChIP assays with GC cell lines and GES-1 cells revealed epigenetic modifications (H3K9me3 and H3K27me3) in the promoter of Cul9 promoter. Data are shown as mean ± SEM; **P < 0.01; ***P < 0.001.

Since epigenetic modification downregulated the levels of tumor-suppressor genes [25], we then analyzed CpGs residing within the *Cul9* promoter using GC TCGA samples. It was observed that the promotor of *Cul9* was hypermethylated in GC tumors rather than in normal controls (Figure 1E). And CpG methylation sites were negatively correlated with *Cul9* expression (Figure 1F). Importantly, CpG methylation of *Cul9* could predict worse survival rates in GC patients (Figure 1G). Consistent with these bioinformatics data, RT-PCR and ChIP assays with GC cell lines and patients’ samples revealed the epigenetic methylation modifications of the *Cul9* promoter (Figure 1Hi-Hiv). Together, methylation-associated downregulation of Cul9 levels may promote the aggressive progression of GC.

### Cul9 suppresses the proliferation of GC cells *in vitro* and *in vivo*

Next, we tried to reveal the functional roles of Cul9 in GC. CCK-8 assays indicated that knocking down *Cul9* contributed to the growth of BGC823 and HGC27 cells, but the inhibitory effect was observed in *Cul9-*overexpressed GC cells (Figure 2A). EdU assays further demonstrated a tumor-suppressor function of *Cul9* in GC cells (Figure 2B-E). Not surprisingly, reintroducing Cul9 rescued the phenotype induced by shRNA against *Cul9* in BGC823 and HGC27 cells (Figure 2F).

**Figure 2.**
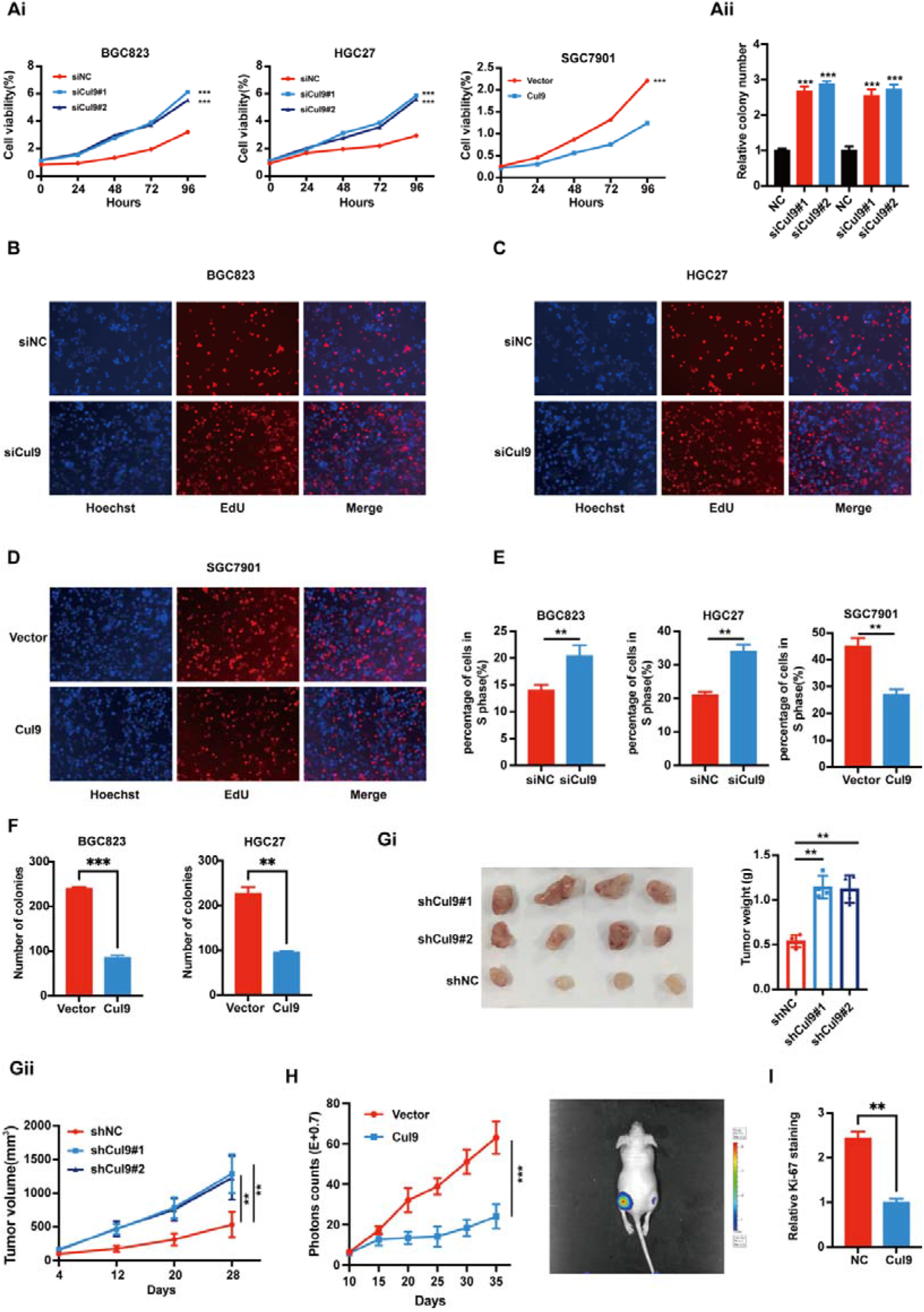
Decreasing Cul9 drives the development of GC. (A) CCK8 assays and colony formation experiments revealed the roles of *Cul9* in regulating the growth of GC cells, respectively. (B-E) Edu assays in three GC cells treated as in (A). The representative pictures were shown. (F) The rescue experiments of *Cul9* in both BGC823 and HGC27 cells with *Cul9* knockdown. (G-I) Xenograft experiments (n=4/group) were performed with GC cells by knocking down or introducing *Cul9*, respectively. Tumor sizes and weights or representative images were presented. Data are shown as mean ± SEM; **P < 0.01; ***P < 0.001.

Subcutaneous tumor experiments in nude mice also were performed. Knocking down *Cul9* led to a significant increase in tumor volume and weight in xenografted mice (Figure 2Gi-ii). However, *Cul9* overexpression inhibited GC development (Figure 2H). The levels of Ki67, a marker of proliferation, were also observed to be negatively associated with the elevated Cul9 (Figure 2I). Therefore, we conclude that Cul9 blocks the growth of GC *in vitro* and *in vivo*.

### Silencing Yes1 at least partially counteracts the growth of GC cells enhanced by Cul9 deficiency

To search for the possible effector downstream of Cul9, three experiments were carried out as follows. First, the GC TCGA datasets were divided into two groups: *Cul9* ^high^ and *Cul9* ^low^ (Figure 3A). Function pathways enriched by the top 500 genes in either group suggest that the levels of *Cul9* expression are mainly associated with tyrosine kinase, MAPK kinase, and nucleotide metabolism pathways, respectively (Figure 3B and Supplementary Table S1). Based on these results, we tried to clarify which tyrosine kinase is associated with Cul9. Given that Cul9 is an ubiquitination E3 ligase, we expected that Cul9 deficiency may upregulate this tyrosine kinase. Thus, we knocked down *Cul9* in BGC823 and HGC27 cell lines. MS/MS analysis revealed that Yes1 is only one tyrosine kinase among the up-regulated 22 proteins according to the overlaps (Figure 3C). To further confirm this result, GC cells with *Cul9* loss were treated with or without a panel of 416 kinase inhibitors. This kinase inhibitor library screen indicated that CH6953755, a Yes1 inhibitor, was most active to block the cell viability and colony formation induced by *Cul9* loss (Figure 3D-Ei-iv). Yes1 knockdown also presented similar effects as CH6953755 (Figure 3F). Therefore, *Cul9* loss promotes GC growth in a Yes1-dependent manner.

**Figure 3.**
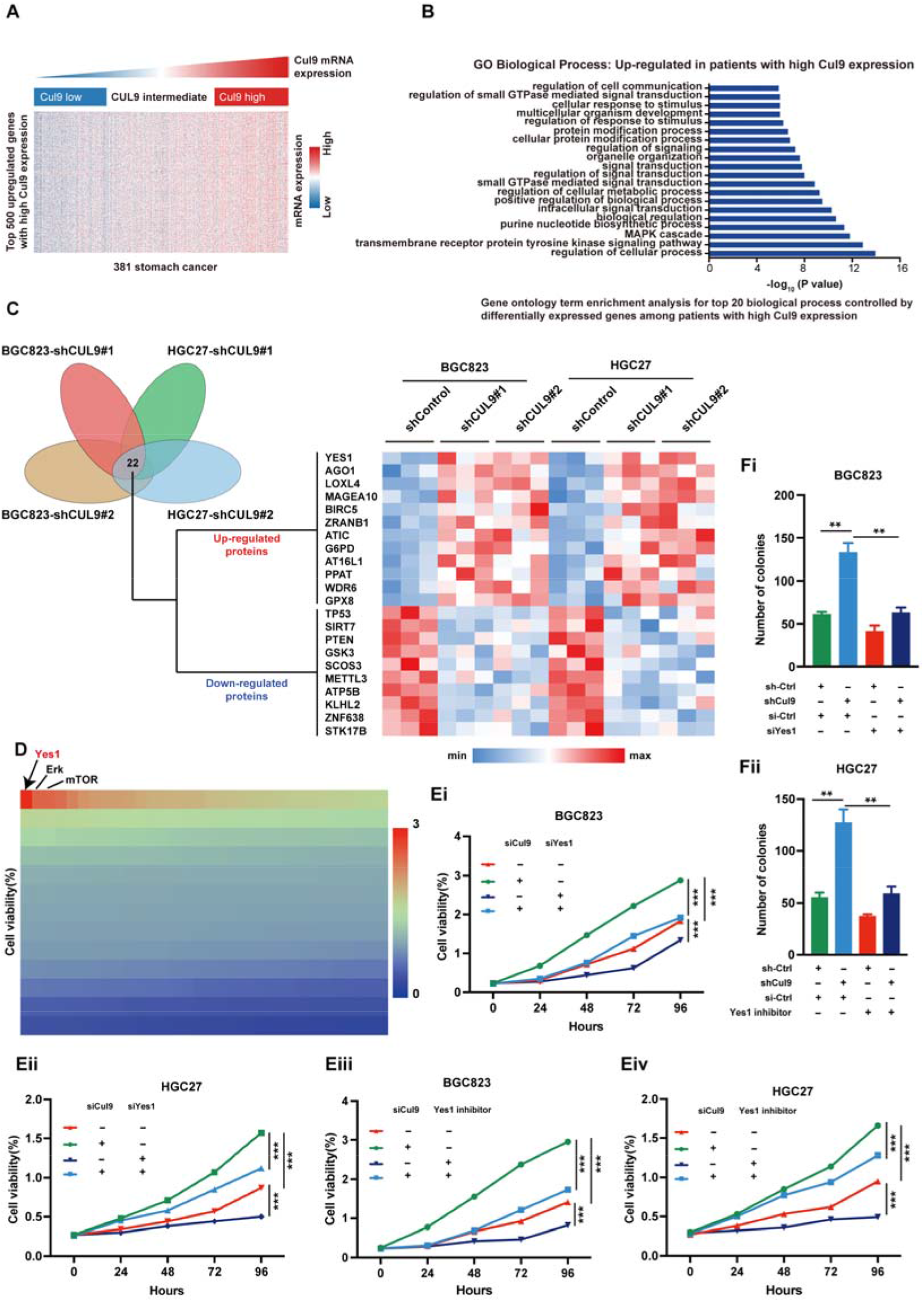
Silencing Yes1 at least partially counteracts the growth of GC cells enhanced by Cul9 deficiency. (A) Heatmap showing the effects of *Cul9* levels on the transcriptomics in GC TCGA database. (B) The enrichment of top 20 function pathways modulated by *Cul9* expression in GC patients. (C) *Cul9* was knocked down in BGC823 and HGC27 cell lines. Mass spectrometry was performed. The Venn diagram indicated the overlap of proteins differentially expressed in BGC823 and HGC27 cells transfected with *shCul9-1*# or *2*#. And the heatmap of the upregulated and downregulated proteins was generated to predict potential Cul9-associated tyrosine kinase candidates. Finally, Yes1 was selected according to the overlaps. (D) BGC823 and HGC27 cells with *Cul9* knockdown were treated with or without 10 mmol/l 416 kinase inhibitors, and cell viability was analyzed. The relative effects of these inhibitors were presented using the heat map. The top three inhibitors having anti-GC activity were listed. (Ei–iv) BGC823 and HGC27 cells with *Cul9* knockdown were treated with or without 10 mmol/l Yes1 inhibitor CH6953755. Indicated experiments were performed (n=3, *P < 0.05). (F) BGC823 and HGC27 cells with *Cul9* knockdown were transfected with Yes1 siRNA for 48 hours. Then examined the cell viability. Data are shown as mean ± SEM; **P < 0.01; ***P < 0.001.

### The E3 ligase activity of Cul9 is required for Yes1 degradation

Next, we tried to elucidate how Cul9 regulated Yes1 in GC. First, we observed that the protein levels of Yes1 in *Cul9*^−/−^ MEFs were increased compared with those in *Cul9*^+/+^ MEFs (Figure 4A). Consistently, overexpression of Cul9 decreased the protein levels of Yes1 in GC cells (Figure 4B). Although we observed an accumulation of Yes1 proteins in siRNA-*Cul9* GC cells, this did not upregulate the mRNA levels of Yes1 (Figure 4C). CHX experiments indicated that the protein half-life of Yes1 was longer in the *Cul9^-/-^* MEFs than in the *Cul9*^+/+^ MEFs (Figure 4D), while the addition of MG132 stabilized the protein levels of Cul9 in both cell types (data not shown).

**Figure 4.**
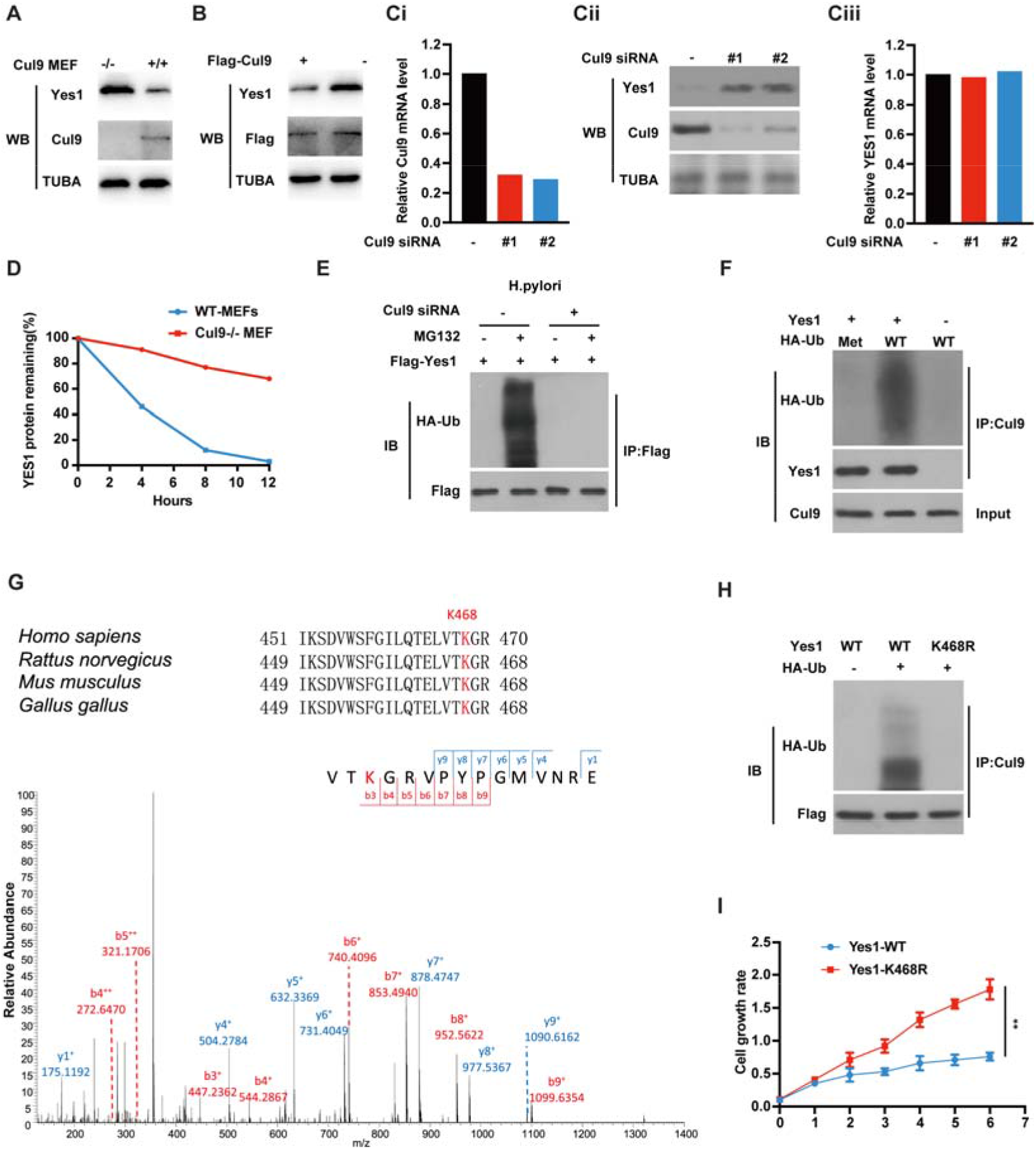
The ligase activity of Cul9 is necessary for Yes1 degradation. (A) The protein levels of Yes1 were examined in *Cul9*^+/+^ and *Cul9*^−/−^MEFs. (B) Endogenous Yes1 levels were determined in GC cells transfected with *Flag-Cul9* by western analysis using indicated antibodies. (C) *Cul9* was knocked down in SGC7901 cells with two siRNAs. The protein and mRNA levels of Yes1 were examined by Western blots and RT-qPCR, respectively. (D) The cycloheximide-chase experiments were carried out to show the half-life of Yes1 protein in *Cul9*^+/+^ and *cul9*^−/−^MEFs. (E) Yes1 ubiquitination *in vivo* was analyzed using the immunoprecipitates from HGC27 cells which were transfected with scramble or *siCul9*, *HA-ubiquitin, Flag–Yes1*, and then treated with *H.pylori* or MG132 together. (F) *In vitro* ubiquitination assay. The immunoprecipitated Cul9, recombinant Yes1, WT, or methylated (Met) ubiquitin (a negative control). (G) MS/MS identified K468 as a Yes1 ubiquitination site by Cul9, and K468 was conserved residue in Yes1. (H) *In vitro* ubiquitination assay. The immunoprecipitated Cul9 combined with WT-Yes1, or K468R mutant, and recombinant HA-ubiquitin were used. (I) CCK8 assays indicated that the K468R mutant of Yes1 promoted SGC7901 cell proliferation compared with Yes1-WT. Data are shown as mean ± SEM; **P < 0.01.

Given that Cul9 is an E3 ligase, we examined the ubiquitination of Yes1 in *Cul9*-silenced SGC7901 cells. It was observed that the ubiquitination of Yes1 was markedly reduced (Figure 4E). *In vitro*, recombinant Yes1 was ubiquitinated by endogenous Cul9 immunoprecipitated from HGC27 cells (Figure 4F). Furthermore, MS/MS and *in vitro* ubiquitination assays confirm that K468 of Yes1 is the primary site ubiquitinated by Cul9 (Figure 4G-H). Notably, the K468R mutant of Yes1 promoted SGC7901 cell proliferation compared with Yes1-WT (Figure 4I). Together, these findings suggest that Cul9 inhibits GC by ubiquitinatingYes1.

### High-throughput screen identifies HG78 compound as an activator of MBD2 which inhibits the methylation of cul9 promoter and GC cell proliferation

It has been well demonstrated that several methyl-CpG–binding domain (MBD) proteins can read and modulate DNA methylation [26]. These methylation readers are MBD1, MBD2, MBD3, MBD4, and MeCP2 [26]. We wondered which methylation readers could regulate the levels of Cul9 through DNA demethylation. MethPrimer website has predicted the CpG islands in the *Cul9* promoter (Figure 5A). ChIP assays showed that it was MBD2 rather than other readers to directly bind to the promoter of *Cul9* (Figure 5B). To elucidate the effect of MBD2 on the Cul9, we first cloned the methylation sequences in the *Cul9* promoter into a *pCpGI* luciferase construct without CpG, and then transfected this *pCpGI* reporter with inactive mutated *MBD2 (mtMBD2*) or *WT-MBD2* into HEK293 cells, respectively. It was observed that *WT-MBD2*, but not *mtMBD2,* promoted Cul9 expression (Figure 5C). Furthermore, the pCpGI with Cul9 methylation could be inhibited by endogenous MBD2-bound DNA and ectopic MBD2 (Figure 5D). Consistently, MBD2 overexpression increased the levels of Cul9 while knocking down MBD2 had the opposite effect (Figure 5E-F). Therefore, MBD2 demethylates the promoter of Cul9 and is an activator of Cul9.

**Figure 5.**
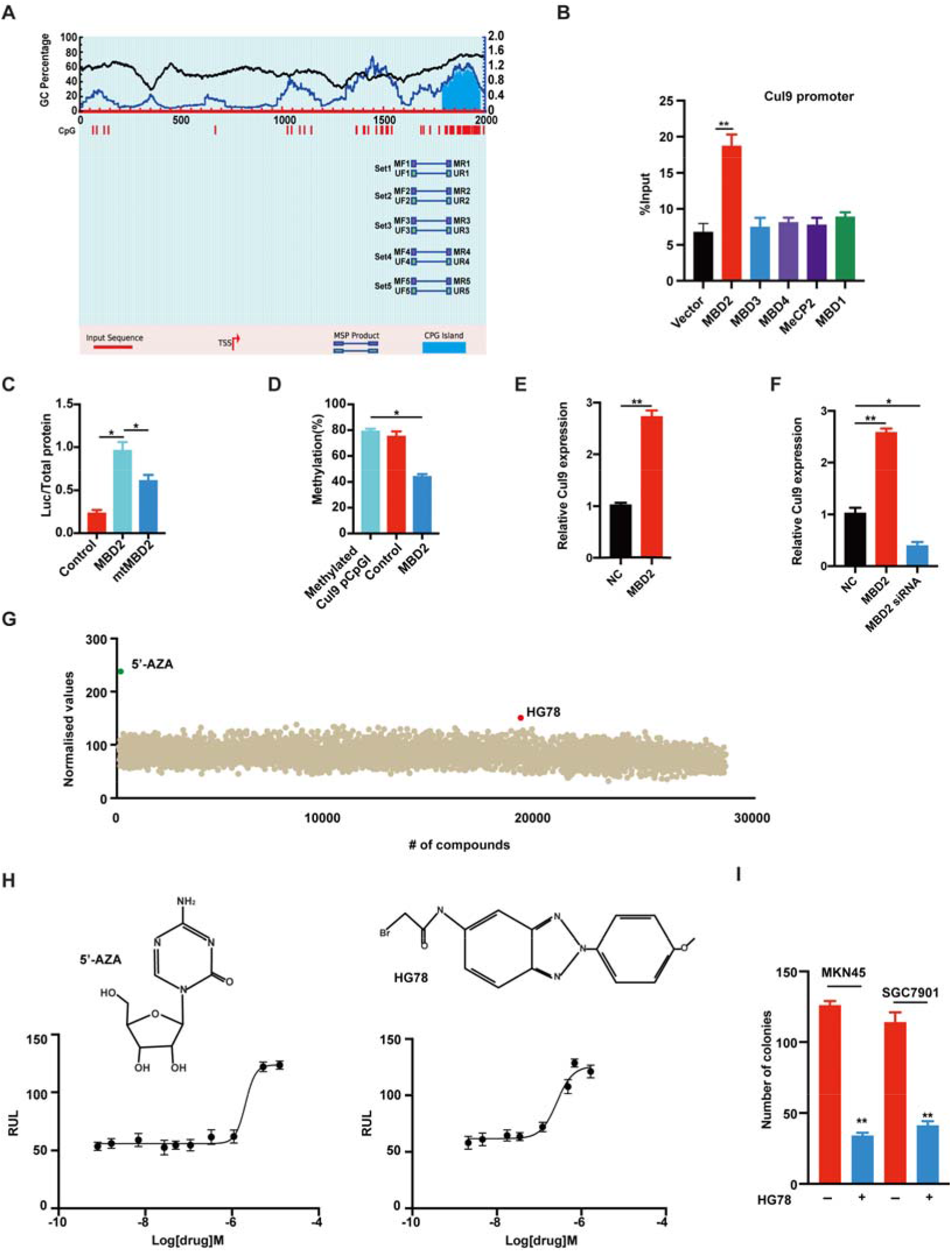
High-throughput screen identifies HG78 compound as an activator of MBD2 which inhibits the methylation of cul9 promoter and GC cell proliferation. (A) Five pairs of primers AND CpG islands were shown in the promoter region of *Cul9*. (B) ChIP experiments using normal gastric cells indicated the direct binding of MBD2 to the promoter of *Cul9*. (C) The effects of MBD2 or mtMBD2 on *Cul9* pCpGI methylation in GC cells. (D) The methylation assays in the CpG islands of *Cul9*. (E) Cul9 levels were examined as indicated. (F) Cul9 levels were presented with a Greyscale image. (G) The dot plot indicated a luminescence-based screen, and HG78 (in red) was identified as a novel activator of MBD2. Green represented 5-Aza. (H) Upper: the chemical structure of 5-Aza and HG78. Lower: the graph indicating the response of HG78 or 5-Aza dose to the reporter of Cul9 in GC cells (n = 3). (I) The effects of HG78 on GC cells and Cul9 expression. Data are shown as mean ± SEM; *P < 0.05; **P < 0.01.

Based on these findings, we then wished to identify one activator of MBD2 which can inhibit the growth of GC cells with Cul9 dysregulation. To this goal, we screened over 28,000 small-molecule compounds from multiple chemical libraries by employing a MBD2-luciferase reporter cell line. Through this luminescence-based approach, we identified HG78 as a potential *MBD2*-reactivating agent (Figure 5G). We also examined the effect of HG78 on *Cul9*-luciferase reporter in GC cells by using 5-azacytidine (5-Aza) as a positive control. We found that their efficacy in the level of Cul9 upregulation was comparable (*E*_max_ = 120∼125 RLU) with the EC_50_ of 5.79 μM and 0.18 μM for 5-Aza and HG78 compounds, respectively (Figure 5H). Furthermore, we demonstrated that HG78 is capable of inhibiting GC cell proliferation in different cellular contexts (Figure 5I). Thus, reactivating MBD2 by HG78 compound could provide therapeutic intervention for GC cells with Cul9 dysregulation.

### Yes1 is identified as a kinase responsible for Cul9 phosphorylation at Y1505 in GC

To search for the upstream regulator of Cul9 activity in GC, we performed the proximity-dependent biotin (BioID2) experiment [27]. Interestingly, this screening identified Yes1 in Cul9-complex (Figure 6A). Since Yes1 is a tyrosine kinase, we hypothesized that it might affect the activity of Cul9 by phosphorylation. To this end, the binding Cul9 to Yes1 exogenously and endogenously was demonstrated (Figure 6Bi-ii and data not shown). We also performed pull-down experiments. The N terminus of Yes1 containing SH2 domain was required for binding to cul9 whereas Yes1 mainly interacted with the N-terminus of GST-Cul9 (Figure 6C). Proximity ligation assay further confirmed that *Helicobacter pylori* (*H.pylori*) promoted the binding of Cul9 to Yes1 in GC cells which mainly happens in the cytoplasm (Figure 6D).

**Figure 6.**
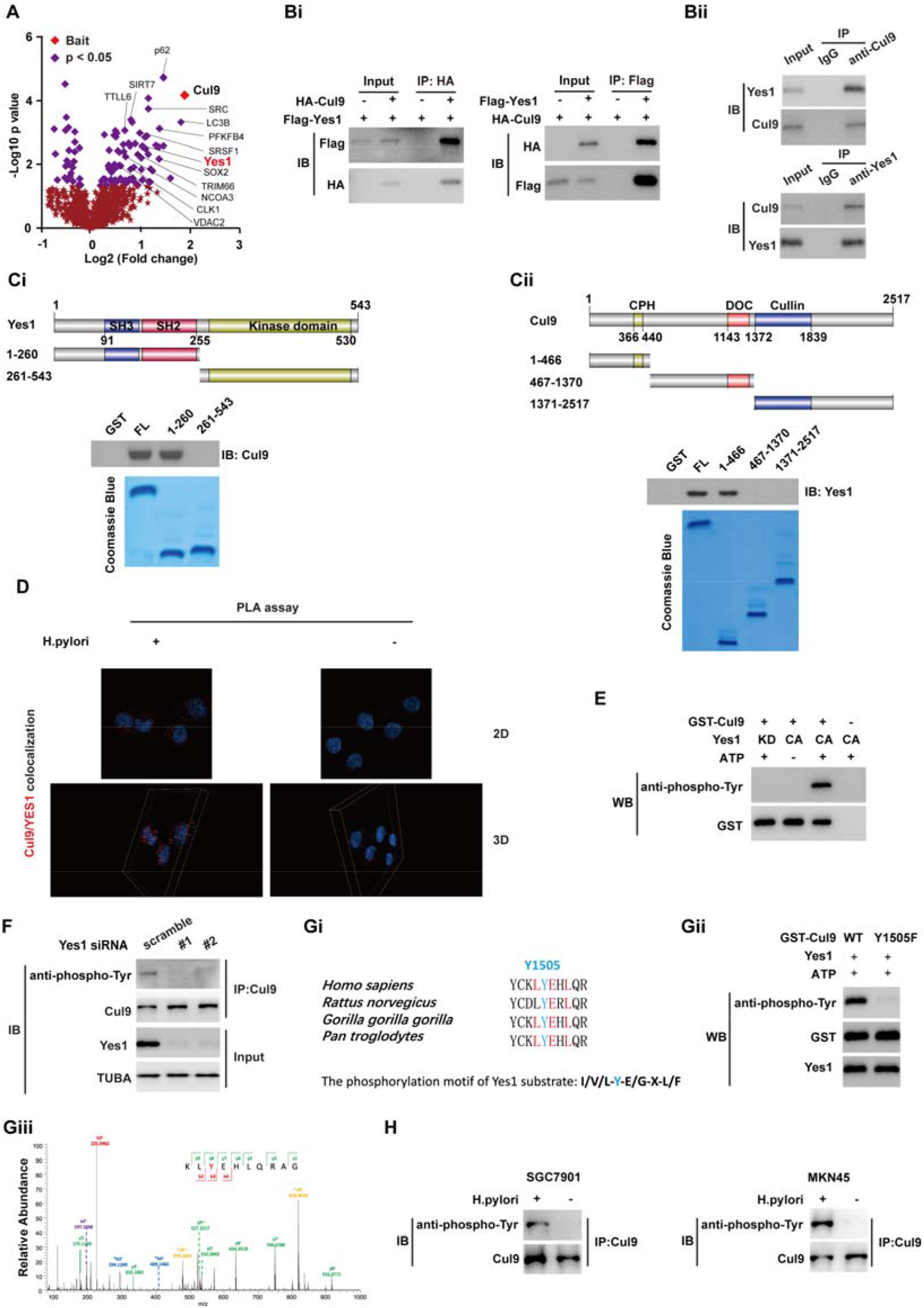
Yes1 is identified as a kinase responsible for cul9 phosphorylation at Y1505 in GC. (A) The proximity-dependent biotin (BioID2) experiment identified Yes1 in Cul9-complex. (Bi) HEK293T cells were co-transfected with HA-Cul9 and Flag-Yes1 for 48 hours. The indicated immunoprecipitation and subsequent immunoblotting were performed with HA or Flag antibodies. (Bii) GC cells were immunoprecipitated with an anti-Yes1 or anti-Cul9 antibody and probed with indicated antibodies. (Ci) Binding of different human Yes1 truncated fragments to Cul9 as indicated (left panel). (Cii) Binding assays about the interaction of Cul9 domains with Yes1 (right panel). The immunoblots and Coomassie blue staining were performed. (D) Proximity ligation assay indicated that *H.pylori* transfection promoted the binding of Cul9 to Yes1 in the cytoplasm of SGC7901 cells. (E) *In vitro* kinase analysis using inactive or active Yes1, ATP, and GST-Cul9. 4G10 antibody detected Cul9 tyrosine phosphorylation. (F) Cell lysates were prepared from SGC7901 cells with Yes1 silencing by using siRNA. Phosphorylation of Cul9 was evaluated by using the indicated antibodies. (Gi and Giii) MS/MS identified Y1505 as a majorly phosphorylated site of Cul9, which is localized in the conserved phosphorylation motif of Yes1. (Gii) *In vitro* Kinase assays were performed with purified GST-Cul9 and its mutant Y1505F. Phosphorylation of Cul9 was evaluated by Western blot with an 4G10 antibody. (H) GC cells were treated with or without *H. pylori* for 6 hours and subjected to immunoblotting analyses with a specific Cul9-Y1505 phosphorylation antibody.

Next, we examined whether Cul9 was an unknown substrate of Yes1 by performing *in vitro* kinase assays. As shown in Figure 6E, Cul9 was phosphorylated in the presence of both ATP and active recombinant Yes1 while an inactive Yes1 mutant lost the ability to phosphorylate Cul9. Consistently, the phosphorylation levels of Cul9 were dramatically reduced when Yes1 was knocked down in SGC7901 cells (Figure 6F). MS/MS identified Y1505 as a majorly phosphorylated site of Cul9, which is localized in the conserved phosphorylation motif of Yes1 [28] (Figure 6Gi and Giii). Furthermore, *in vitro* kinase assays confirmed that Yes1 enhanced phosphorylation of WT-Cul9 rather than its mutant at Y1505F (Figure 6Gii). In addition, by using a specific antibody against phosphorylated Cul9 at Y1505, it was observed that treatment of SGC7901 and MKN45 cells with *H. pylori* significantly enhanced endogenous phosphorylation of Cul9 at Y1505 (Figure 6H). Therefore, we conclude that Yes1 directly binds to Cul9 and phosphorylates it at Y1505 in GC cells.

### Yes1-mediated Y1505 phosphorylation results in selective autophagy of Cul9

Phosphorylation modification is usually linked to protein stability [28]. We then explored whether Yes1-mediated Cul9 phosphorylation at Y1505 affects its stability in GC. As shown in Supplementary Figure 1A, Introducing Yes1 did not reduce the protein expression of Cul9-Y1505F mutant. Since both WT and inactive Yes1 had very little effect on the mRNA levels of either WT-Cul9 or Cul9-Y1505F (Supplementary Figure 1B), implying that Yes1-mediated Cul9 phosphorylation just disrupts its protein stability. Supporting this finding, the half-life of endogenous Cul9 in SGC7901-*shcontrol* cells was dramatically shorter than that in SGC7901-*shYes1* cells (Supplementary Figure 1C-D). Moreover, co-transfection of *Yes1* and *Cul9* did not reduce the half-life of Cul9-Y1505F (Supplementary Figure 1E). Therefore, we conclude that Yes1-mediated cul9 phosphorylation at Y1505 reduces its protein stability.

Since protein degradation is controlled by the lysosome, proteasome, or autophagic pathway [29], we then tried to elucidate the pathway by which Yes1-mediated Cul9 phosphorylation at Y1505 decreases its protein stability. As shown in Supplementary Figure 1F, several autophagy-associated inhibitors such as NH_4_Cl, bafilomycin A1 (BafA1), chloroquine (CQ), and 3-methyladenine (3MA), but not MG132, inhibited Yes1-enhanced degradation of Cul9. Furthermore, this degradation was completely abrogated in the cells with *ATG5* or *BECN1* knockdown (Supplementary Figure 1Gi and Gii). And the addition of BafA1 impeded the degradation of Cul9 in SGC7901 cells (Supplementary Figure 1H). Collectively, Yes1 promotes autophagic degradation of Cul9 protein.

### Yes1 promotes Cul9 binding to p62 which is necessary for selective autophagy of Cul9

Several receptors contribute to selective autophagy [30]. Given that p62, but not NBR1, Nix, NDP52, OPTN, and Tollip, was an interactor of Cul9 (Figure 6A), we then examined its role in the autophagic degradation of Yes1-mediated Cul9. Interestingly, Yes1 only enhanced the bingding of Cul9 to p62 (Supplementary Figure 2A). But *Yes1* deficiency blocked the binding of p62 to endogenous Cul9 (Supplementary Figure 2B). Consistently, Cul9-Y1505F could not form a complex with p62 (Supplementary Figure 2C). Yes1 did not promote Cul9 degradation in cells with *p62* deficiency (Supplementary Figure 2Di-ii). The CHX assays indicated that Cul9 degradation was abolished in *p62* knockdown cells (Supplementary Figure 2E), suggesting that Yes1 facilitates the binding of Cul9 to p62 for its selective autophagic degradation.

Given that selective autophagy required the interaction of p62 to ubiquitinated protein [30], we then examined whether Y1505 phosphorylation induced the ubiquitination of Cul9. As shown in Supplementary Figure 2F, WT-Cul9 rather than Cul9-Y1505F mutant had a K63-linked poly-ubiquitination signal. NH_4_Cl addition induced WT-Cul9 ubiquitination (Supplementary Figure 2G). As expected, adding vanadate also enhanced the levels of Cul9 ubiquitination and pY1505 phosphorylation (Supplementary Figure 2H). Collectively, Yes1-mediated phosphorylation at Y1505 promotes selective autophagy of Cul9.

### COP1 mediated Cul9 K63-linked ubiquitination and selective autophagy

Next, we performed MS/MS to identify the E3 ligase which contributes to the K63-linked ubiquitination of Cul9. This assay revealed two E3 ligases COP1 and SPOP in Cul9-complex. However, co-transfection experiments indicated that overexpression of COP1 rather than SPOP induced K63-linked ubiquitination of Cul9 (Supplementary Figure 3A). On the contrary, silencing *COP1* reduced Cul9 ubiquitination (Supplementary Figure 3B). Furthermore, the E3 ligase-defective COP1-C156/159S mutant did not induce Cul9 K63-linked ubiquitination (Supplementary Figure 3A).

Given that the VPD/E motif in the target protein is important for COP1 binding [31], the VP residues of Cul9 protein located in (aa 2461-2480) were mutated with “AA”. It was found that COP1 could not interact with the Cul9-VP/AA mutant (Supplementary Figure 3Ci-ii).

We also demonstrated that the binding of Cul9 to COP1 required Yes1-mediated Y1505 phosphorylation (Supplementary Figure 3Di-ii).

MS/MS experiments further identified the K1657 in Cul9 as a major ubiquitination site modified by COP1 (Supplementary Figure 3E-F). Not surprisingly, autophagic receptor p62 did not interact with the Cul9-K1657R mutant (Supplementary Figure 3G), which is consistent with the previous data.

### GC-related mutants of Yes1 or *H. pylori* markedly promote Y1505 phosphorylation of Cul9

Given that the carcinogenesis of GC was closely associated with *H. pylori* (HP) infection [14], we examined whether *H. pylori* infection regulated the activity of Yes1 and Cul9 phosphorylation at Y1505.

First, SGC7901 cells were infected with J166 and 7.13 two *Cag^+^ HP* strains, respectively, and the levels of Yes1 protein rather than mRNA were significantly upregulated (Figure 7A-Bi). Interestingly, when these cells were incubated with BafA1, Cul9 K63-linked poly-ubiquitination and phosphorylation at Y1505 were upregulated, and binding of p62 to Cul9 was observed (Figure 7Bi). However, in the cells without BafA1 incubation, *HP* infection leads to a reduction in Cul9 protein levels (Figure 7Bii). Not surprisingly, *HP* infection significantly enhanced the activity of Yes1 in GC (Figure 7C).

**Figure 7.**
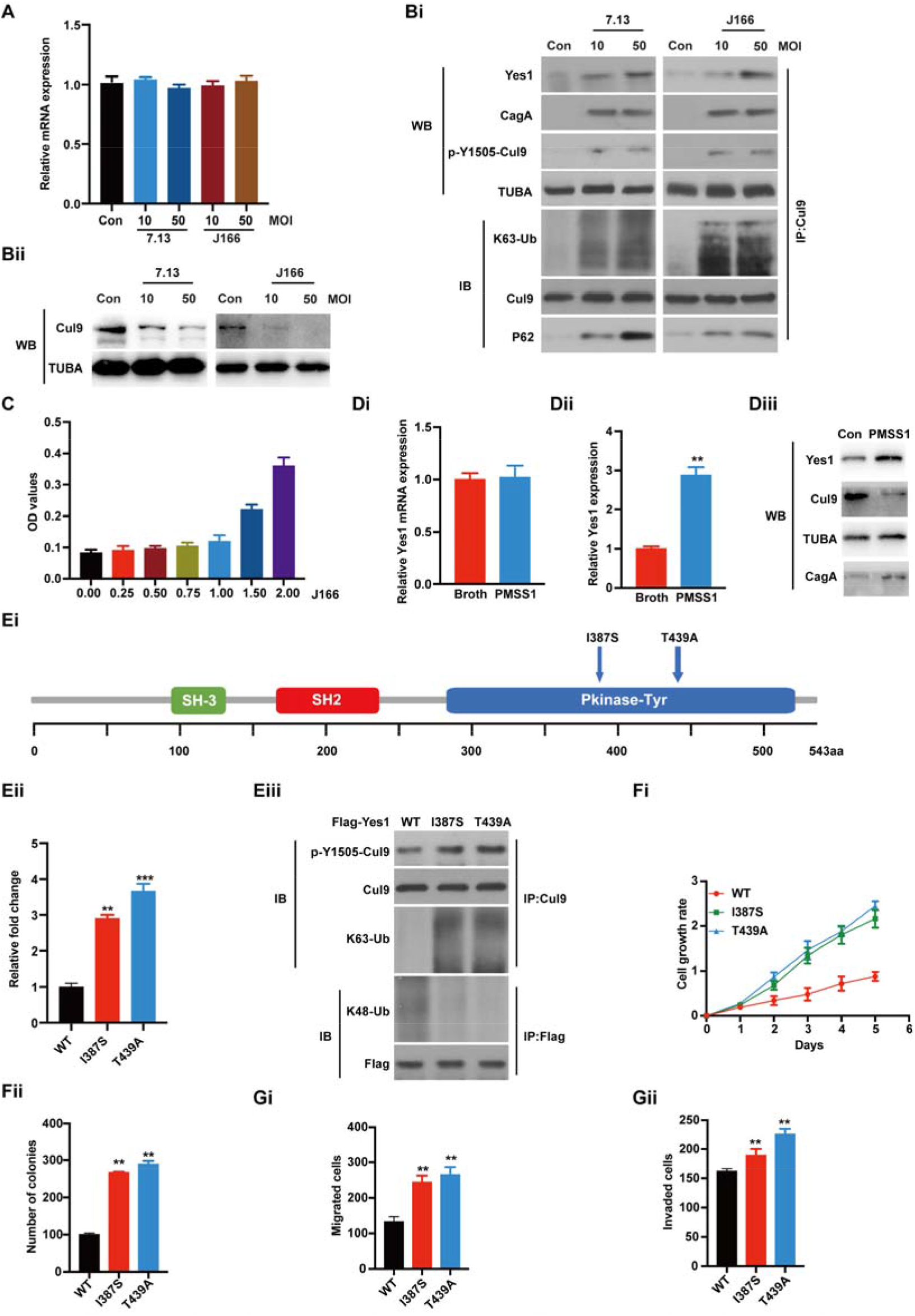
GC-related mutants of Yes1 or *H. pylori* markedly promote Y1505 phosphorylation of Cul9. (A) The levels of Yes1 in SGC7901 cells were challenged with or without *CagA+ HP* strains 7.13 or J166. MOI means a multiplicity of infections. (Bi) Western blot assays for the samples from (A) treated with BafA1. (Bii) SGC7901 cells infected with 7.13 and J166. The experiments were performed as indicated. (C) Yes1 activity was analyzed. (D) Mice were challenged with *HP* strain PMSS1 or Brucella broth as a control. QRT-PCR (Di) and Western blots (Dii-iii) were performed. (Ei) Yes1 architecture domain and its GC-related mutants in GC patients. (Eii) Relative activity of Yes1 was analyzed. (Eiii) IP kinase assay. Briefly, Yes1 was immunoprecipitated from HEK293 cells expressing Flag-tagged WT-Yes1 or its mutants. Then the immunoprecipitated Yes1 was mixed with [^32^P] ATP and a known substrate peptide of Yes1. The results were normalized to 1.0 for WT-Yes1. (Fi-ii) The effects of GC-related Yes1 mutants on GC cells. (Gi-ii) The invasion and migration of GC cells were affected by GC-associated Yes1 mutants. Data are shown as mean ± SEM; **P < 0.01; ***P < 0.001.

*In vivo*, in mice infected with PMSS1-*HP* strain, the expression of Yes1 protein rather than mRNA was increased, while this effect was not observed in mice treated with Broth (as a negative control) (Figure 7Di-ii). Moreover, challenging *HP* markedly downregulated Cul9 levels in mice (Figure 7Diii).

Clinically, two somatic missense Yes1 mutations (I387S and T439A) were previously revealed in the GC-TCGA database (Figure 7Ei), but their functional significance was not elucidated. As shown in Figure 7Eii-iii, I387S and T439A mutants in the kinase domain greatly increased the activity of Yes1 but decreased its K48-linked poly-ubiquitination. Consistently, I387S and T439A mutants enhanced Cul9 K63-linked poly-ubiquitination and phosphorylation at Y1505 (Figure 7Eiii). Importantly, GC cells with I387S and T439A mutants were more aggressive than the controls (Figure 7Fi-ii and Gi-ii).

Together, GC-associated Yes1 mutants and *HP* infection enhance Yes1 activity, upregulated Cul9 K63-linked poly-ubiquitination, and Y1505 phosphorylation.

### *Knockin* of *Cul9-Y1505D mutant* in *gp130*^F/F^ mice confers an overall metabolic advantage, thereby promoting gastric tumorigenesis

To interrogate the role of Yes1-mediated Cul9 phosphorylation at Y1505 in GC, *gp130*^F/F^ mice with *WT-Cul9* and *Cul9-Y1505D knockin* were generated (Figure 8A-C). At 11 weeks of age, we did not observe an obvious difference in the tumor mass and stomach size between *gp130*^F/F^; *WT-Cul9* and *gp130*^F/F^; *Cul9-Y1505D* mice. However, 20-week-old *gp130*^F/F^; *WT-Cul9* mice stomachs displayed a smaller tumor mass and incidence (Figure 8Di-ii), which corresponded with smaller hyperplastic lesions in *gp130*^F/F^; *WT-Cul9* mice (Figure 8E). In addition, *gp130*^F/F^; *WT-Cul9* mice had a better percent survival than *gp130*^F/F^; *Cul9-Y1505D* mice at the corresponding ages (Figure 8F). These data, therefore, confirm that Yes1-mediated Cul9 phosphorylation at Y1505 promotes gastric tumorigenesis in *gp130*^F/F^ mice.

**Figure 8.**
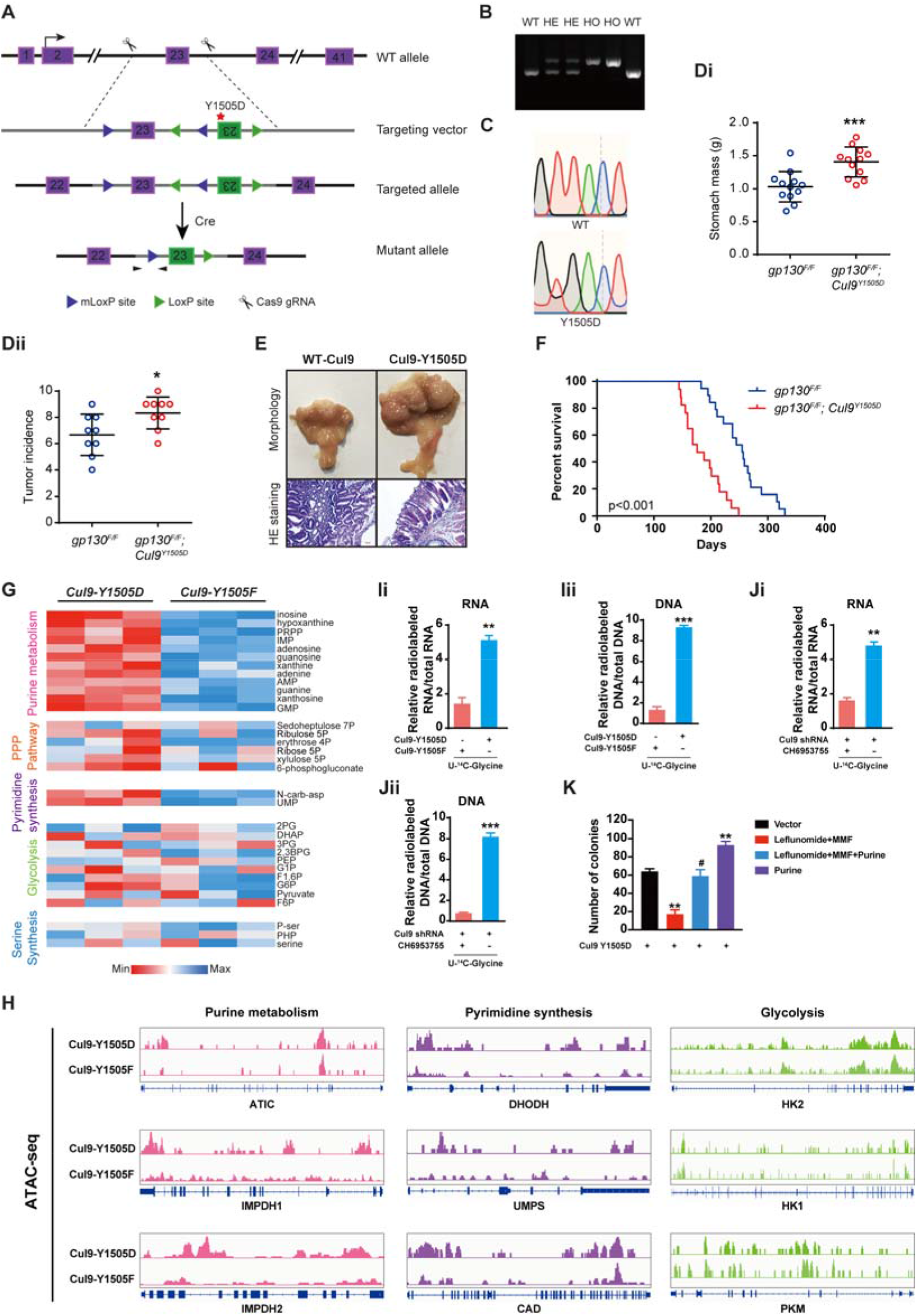
*Knockin* of *Cul9-Y1505D mutant* in *gp130*^F/F^ mice confers an overall metabolic advantage, thereby promoting gastric tumorigenesis. (A) Establishing *WT-Cul9* and *Cul9-Y1505D* knockin mice. Schematic genome maps of wild-type and targeted alleles were shown. Loxp sites were shown. (B) Representative genotyping PCR result. (C) Sequencing confirmed mutant site. (D) Scatter plots showing the mass (Di) of mice stomachs and gastric tumors, and the incidence (Dii) of tumors at 20-week-old age. (E) Representative H&E-staining and tumor pictures at 20-week-old age. (F) Mice survival curve. (G) The metabolic profiling in SGC7901 cells as indicated. (H) ATAC sequence of SGC7901 cells as indicated. (I) *Cul9-Y1505D* and *Cul9-Y1505F* mutants differently affected RNA and DNA biosynthesis in SGC7901 cells, respectively. (J) The Yes1 inhibitor rescued the phenotype of Cul9 loss in SGC7901 cells. (K) Leflunomide and Mycophenolate mofetil (MMF), two inhibitors targeting purine and pyrimidine synthesis pathways, respectively, inhibited the growth of SGC7901 cells expressing Cul9-Y1505D. Data are shown as mean ± SEM; *P < 0.05; **P < 0.01; ***P < 0.001.

Since metabolism reprogramming dictates the development of GC, we then examined whether Yes1-mediated Cul9-Y1505 phosphorylation reprogrammed GC metabolism. To this end, *Cul9-Y1505F* or *Cul9-Y1505D* mutant was reintroduced into SGC7901 cells with stable knockdown of endogenous Cul9. The metabolic profiling indicated that SGC7901 cells expressing *Cul9-Y1505D* rather than *Cul9-Y1505F* mutant markedly promoted the purine and pyrimidine synthesis pathway (Figure 8G and Supplementary Table S2). These findings were further confirmed by ATAC sequence (Figure 8H). Supporting these findings, Y1505 phosphorylation of Cul9 in SGC7901 cells markedly promoted the biosynthesis of RNA and DNA, as compared to the cells expressing Cul9-Y1505F (Figure 8Ii-ii). Notably, the Yes1 inhibitor rescued the phenotype of Cul9 loss (Figure 8Ji-ii), further confirming the roles of Cul9-Y1505 phosphorylation in GC metabolism. Importantly, Leflunomide and Mycophenolate mofetil (MMF) targeting purine and pyrimidine synthesis pathways inhibited the proliferation of GC cells with Cul9 loss (Figure 8K). Therefore, Cul9-Y1505 phosphorylation confers marginal metabolic benefits to maximize the growth of GC cells in a Yes1-dependent manner.

### The levels of Yes1 and Cul9 are oppositely related in patients with GC

Finally, we tried to clarify the clinical significance of the findings as described above. It was observed that the levels of Yes1 and Cul9 are oppositely related in patients with GC, while positively linked to pY1505-Cul9, ATIC (purine metabolism), and DHODH (pyrimidine synthesis) (Supplementary Figure 4A–B). Furthermore, GC Patients with lower Cul9 and higher Yes1 had a poor prognosis (Supplementary Figure 4C, Supplementary Table S3 and 4). Therefore, all clinical findings support a novel model about the role of the Yes1-Cul9 crosstalk in GC development (Supplementary Figure 4 D).

## Discussion

Tyrosine phosphorylation is an important research topic because of its critical effect on the functional switch of tumor-associate genes [32]. In this study, we indicate that: 1), *Cul9* promoter is hypermethylated and its levels are markedly reduced in GC cell lines and clinical samples, which predicts a poor prognosis; 2), the tyrosine kinase Yes1 is identified as a novel regulator for cul9 activity and protein stability by directly phosphorylating Cul9 at Y1505, which promotes its selective autophagy degradation; 3), Cul9 in turn competes with Cul7 to bind Yes1 and disrupts Yes1 stability; 4), GC-related Yes1 mutation or *HP* infection induces Cul9-Y1505 phosphorylation and its oncogenic character; 5), Mice with *Cul9-Y1505D knockin* are more susceptible to gastric tumorigenesis compared to WT-Cul9 counterparts; 6), Metabolic profiling indicates that Cul9-Y1505D mutant mainly promotes pyrimidine and purine synthesis pathways in GC, suggesting a nucleotide metabolic reprogramming; 7), The DNA-demethylating drug decitabine or methylation reader MBD2 upregulates Cul9 expression and limits GC cell proliferation in a Yes1-dependent manner; 8), The Yes1 inhibitor CH6953755 or Leflunomide and Mycophenolate mofetil (MMF) suppressing nucleotide metabolism impairs the malignancy of GC with Cul9 dysregulation; 9), High-throughput screen identifies HG78 compound to upregulate Cul9 expression and limits GC cell proliferation by activating MBD2, a CpG methylation reader; 10), Clinically, a significant correlation is observed among high Cul9-phosph-Y1505 levels, Yes1 activity, and nucleotide metabolic markers in different pathological features of GCs. Together, this project highlights an unknown role of the Yes1-Cul9 loop in GC progression, suggesting novel potential therapeutic targets.

An important discovery in this study is the key role of Cul9 in nucleic acid synthesis and GC development. So far, very little was known about the effect of an E3 ligase in purine and pyrimidine synthesis. The previous studies about nucleotide metabolism majorly focused on several transcriptional factors and kinases. For instance, DYRK3 and UHMK1 kinases modulate purine synthesis in hepatocellular carcinoma and GC by controlling the chromatin accessibility of ATF4, respectively [23, 33]. CDC-like kinase 4 (CLK) family kinases oppositely regulate the transcription of purine metabolic genes in cholangiocarcinoma and esophageal carcinoma by regulating c-Myc and MITF, respectively [25, 34]. ERK2 promotes nucleotide metabolism in Kras-related tumors [35]. Herein, the Yes1-Cul9 loop activates both pyrimidine metabolism and purine synthesis pathway. However, the underlying mechanism by which Yes1-mediated Cul9 phosphorylation at Y1505 sustains nucleic acid synthesis remains unclear. Previous studies identified mTORC1 as a key kinase activating both pyrimidine metabolism and purine synthesis pathway [24, 36]. A previous study also indicated crosstalk between Yes1 and mTORC1 [37]. In addition, the activity of Cul7 (a Cul9 partner) is dependent on mTORC1 [38]. All these reports seem to support a possibility: mTORC1 may be downstream of the Yes1-Cul9 loop in metabolic reprogramming. Further study is needed to validate this idea, which will be help explore the potential therapeutic benefit of treating GCs. Notably, when we are performing this manuscript, another group just reported the role of ubiquitin protein ligase E3 component N-recognin 7 (UBR7) in regulating nucleotide biosynthesis in T cell acute lymphoblastic leukemia (T-ALL) [39]. We believe that future studies will identify other E3 ligases involved in purine and pyrimidine metabolism beyond Cul9 and UBR7.

A large number of malignancies are linked to the enhanced activity of SFKs which happens via different mechanisms [40]. Substantial evidence now bolsters the proposal that losing negative regulation such as loss of ubiquitination promotes SFKs deregulation and carcinogenesis [40]. Apart from Cul9 E3 ligase in this study, Cbl-family E3 ligases, Nedd, Cullin-5, SIAH, Nrdp1, ITCH, and CHIP proteins have been reported as negative regulators of tyrosine kinases or SFKs [40]. For example, Cbl-family E3s have been shown to degrade Yes1, c-Src, and Lyn by ubiquitination, thereby inhibiting tumor progression [41]. In turn, the activation of c-Src or Yes1 promotes tyrosine phosphorylation of c-Cbl itself that leads to its proteasomal destruction, indicating a reciprocal regulation between c-Cbl E3 ligase and SFKs [42]. Consistent with this literature, ubiquitination of Yes1 by Cul9 is restrained in Yes1-transformed cells, and by promoting the destruction of Cul9, Yes1 enables metabolic reprogramming, which explains Yes1-Cul9 collaboration in GC oncogenesis. Notably, negative regulation of SFKs or RTK by c-Cbl-family E3s has also been shown to act via mechanisms different from ubiquitin-dependent regulation such as functioning as an adapter [41]. Therefore, we wonder whether the effect of Cul9 on SFK signals may be highly context-dependent. If this is the case, the relative importance of Cul9 protein acting through E3 activity or an adapter mechanism should be fully elucidated.

Besides the major finding linking Yes1 to Cul9-Y1505 phosphorylation was shown to be relevant under *H. pylori* infection conditions and exposure of cultured monolayers of human GC cells to *H. pylori* resulted in increased activity of Yes1, our MS/MS data also report that the tyrosine phosphatase PTPN13 co-immunoprecipitates with Cul9. Given the delicate balance between kinase/phosphatase protein phosphorylation, it is not surprising that we postulate that tyrosine phosphorylation of Cul9 may be altered by PTPN13 once *H. pylori* infection for GC cells was removed. This hypothesis was based on the following reasons: 1), PTPN13 was reported to negatively regulate phosphorylation of JAM-A at Y280 [43]; 2), Even though PTPN13 has a potential pro-oncogenic role in malignancies [44], its tumor-suppressor role has been confirmed in several clinical studies [45]; 3), PTPN13 is involved in the oncogenic activities of the members of the Src family [46]. Therefore, it would be particularly interesting to elucidate the effect of PTPN13 on the Yes1-Cul9 loop because validating this idea will provide new insights into tyrosine phosphatase contributions to GC development and Cul9 activity regulation.

It is well-known that tumor cells often use epigenetic mechanisms to regulate the expression of tumor-associated tumor-suppressor genes [47]. Similar to mutational inactivation or gene copy loss, CpG methylation, H3K27me3, and H3K9me3 of these genes also contribute to tumor development [47]. Therefore, *Cul9* promoter methylation represents a *bona fide* GC-driving event, which directly correlates with lower cul9 expression and bad prognosis in patients with GC. The potential benefit of decitabine treatment in GC cells with Cul9 methylation further confirms this model. Notably, TCGA data reveals *Cul9* methylation in other solid tumors, targeting the Cul9-Yes1 loop may have broader implications.

Although it is still controversial whether MBD2 itself is a demethylase, the correlation between DNA demethylation and high levels of MBD2 has been widely reported in breast cancer as well as in several autoimmune diseases [48]. In general, once the methylated DNA is recognized by MBD2, other remodelers or enzymes can be recruited to form a big complex, thereby resulting in the activation of gene expression [48]. Herein, we provide convincing evidence that activating MBD2 upregulates Cul9 expression, and we also validate MBD2 as a potential target for therapeutic intervention in GC with Cul9 dysregulation. Notably, given that mechanical loading may induce the complex formation of MBD2 with tyrosine kinases such as c-Src, Pyk2, or FAK in the nucleus that functions to suppress anabolic gene expression [49], we wonder whether MBD2 can directly bind to Yes1 as a mechanism to prevent the proliferation reaction of GC cells in response to *H.pylori* stimulation. So far, none of the existing drugs has been shown to activate MBD2 and applied to preclinical or clinical trials. Thus, there is a need to find a MBD2-reactivating compound using a library screen. Herein, we tried to identify one small molecular, HG78, that can reactivate MBD2 and alleviate the GC burden. But more studies are needed to fully elucidate the mechanistic basis by which HG78 reactivates MBD2 in GC cells, and whether similar effects can be observed in other human tumors.

In conclusion, the findings in this report are consistent with a model depicted in Supplementary Figure 4D. In this model, the activation of Yes1 by *H. pylori* during inflammation promotes tyrosine phosphorylation of Cul9 at Y1505 and its instability, which results in increased nucleotide metabolism and GC development. In turn, Cul9 inhibits Yes1 expression and activity. Collectively, this study offers some new insights into how the Yes1-Cul9 feedback loop reprograms GC metabolism under inflammatory conditions.

## ACKNOWLEDGMENTS

This work was supported by grants from the science and technology innovation program of Hunan province (2021 RC4056); National Natural Science Foundation of China (81874063, 82073373). Hunan Province Natural Science Foundation (2020JJ4108). We thank Xin Xu for his technical supports and the helpful suggestions for this manuscript.

## Supplementary Figure legends

**Supplementary Figure 1.**
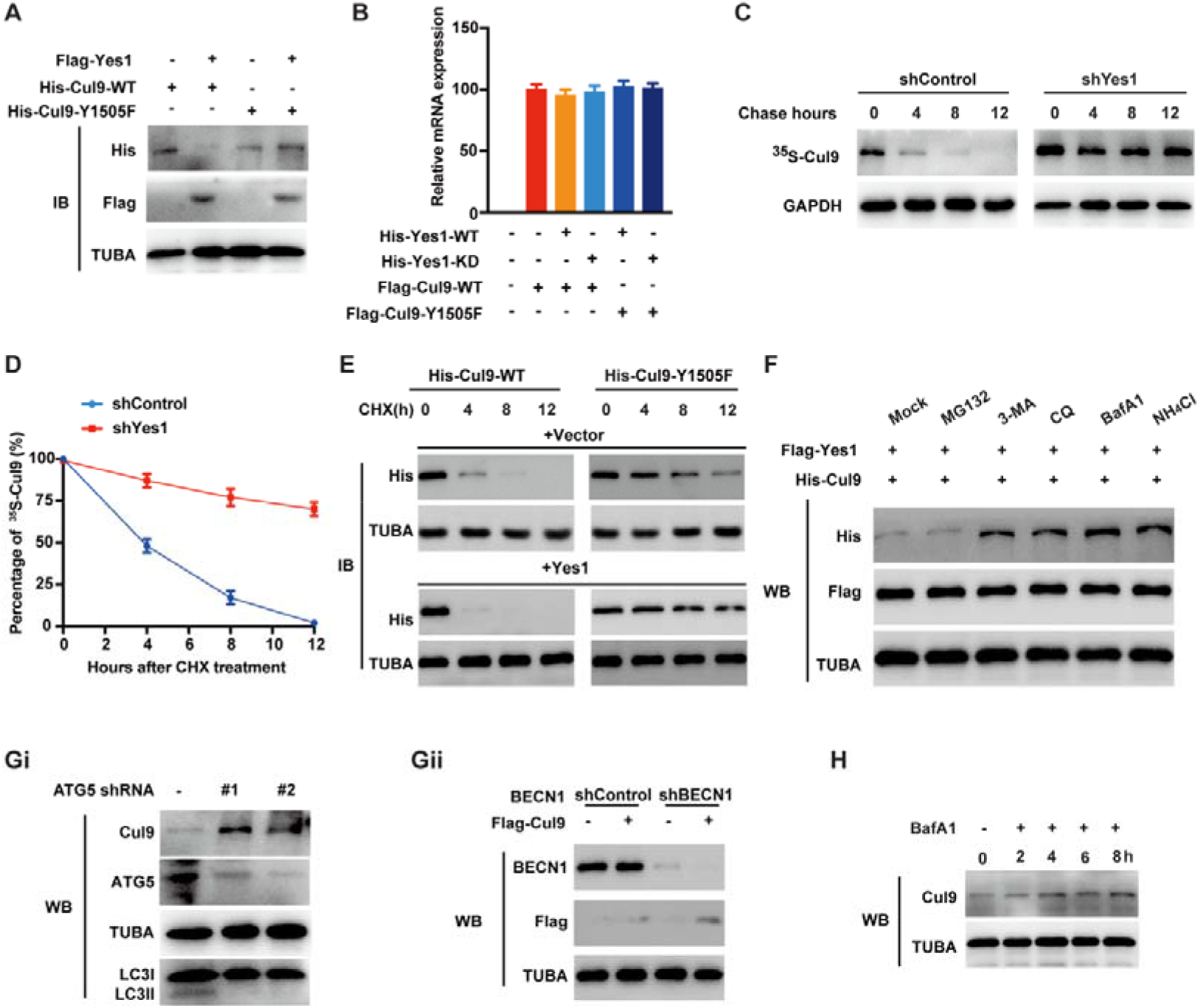
Yes1-mediated Y1505 phosphorylation results in selective autophagy of Cul9. (A) *Flag-Yes1* was transfected into HEK293T cells for 48 hours with or without *His-Cul9*. Western blots were performed as described. (B) HEK293T cells were transfected with *Flag-Cul9* and *WT-His-Yes1* or *Yes1-KD* mutant. The levels of *Cul9* mRNA were examined. (C) [^35^S]-methionine was used to label the cells with SGC7901-*shcontrol* and SGC7901-*shYes1*, and chasing with a standard medium was followed. (D) The endogenous Cul9 expression was examined in (C). (E) The CHX assays were performed. (F) After *Flag-Yes1* and *His-Cul9* co-transfection, HEK293T cells were treated using NH_4_Cl (20mM), 3MA (10mM), MG132 (10μM), CQ (50μM), or BafA1 (0.2μM). Then immunoblots were performed. (Gi and ii) The effects of *Beclin1* or ATG5 knockdown in *Cul9*-transfected HEK293T cells. (H) BafA1 increases Cul9 levels in SGC7901 cells for the duration indicated.

**Supplementary Figure 2.**
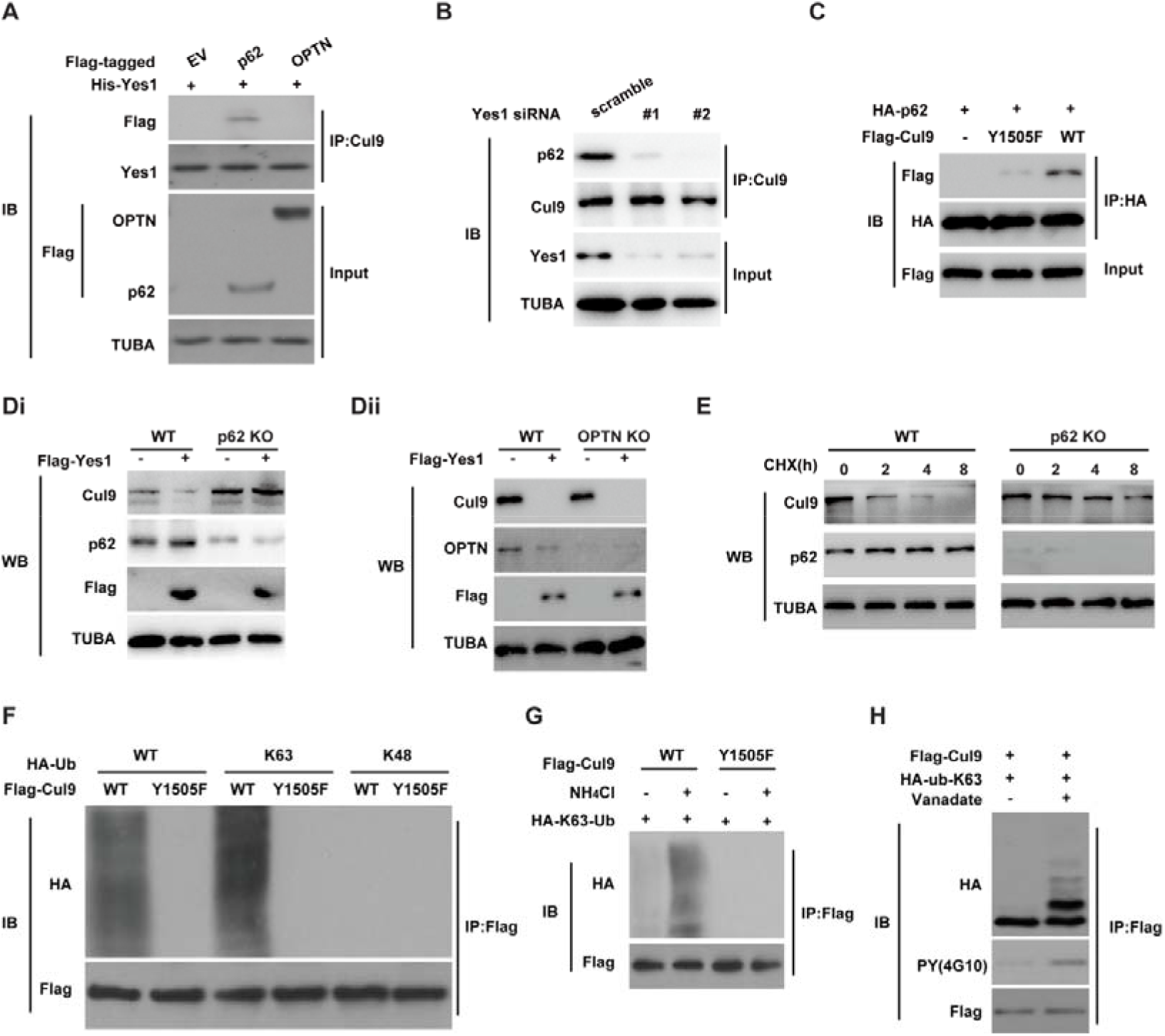
Yes1 promotes Cul9 binding to p62 which is necessary for selective autophagy of Cul9. (A) After *Yes1*, *p62*, and *OPTN transfection*, HEK293T cells were immunoprecipitated with anti-Cul9. Then performing immunoblot analysis as indicated. (B) *Yes1* knockdown disrupted the binding of Cul9 to p62 in SGC7901 cells. (C) HEK293T cells were transfected as indicated, and co-immunoprecipitation was performed. (Di-ii) HEK293T cells with *p62 or OPTN* knockdown or control cells were transfected with *Flag-Yes1*. The immunoblots were performed. (E) Control or *p62*-knockdown HEK293T cells were treated with 100μg/mL CHX as indicated. Then examine the levels of Cul9 protein. (F) Y1505F mutation reduced Cul9-K63 ubiquitination. (G) 15 mM NH_4_Cl treatment increased K63-poly-ubiquitination of Cul9. (H) Vanadate addition enhanced Cul9 ubiquitination.

**Supplementary Figure 3.**
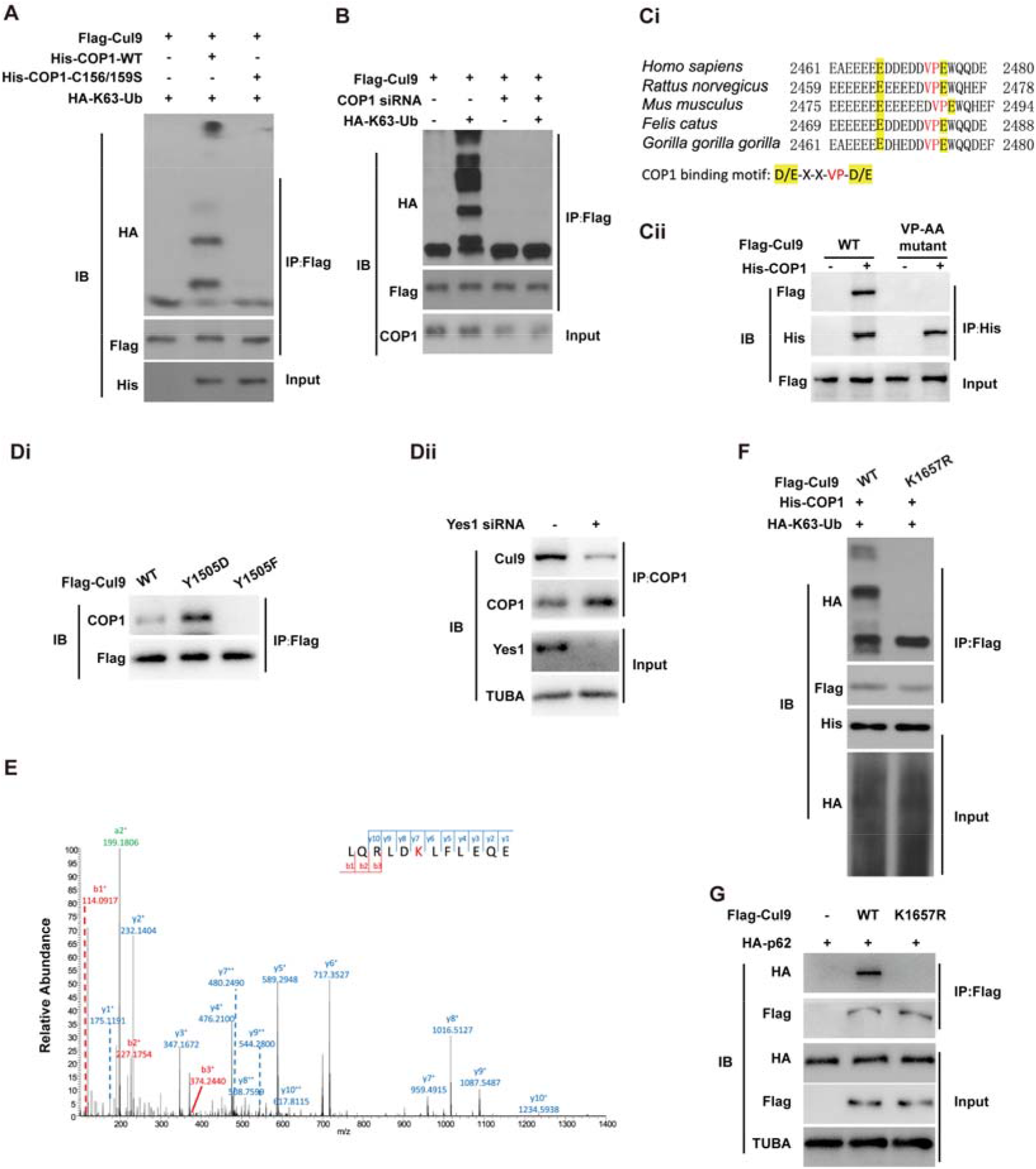
COP1 mediated Cul9 K63-linked ubiquitination and selective autophagy. HEK293T cells were infected using the indicated constructs (A) or scramble or *siCOP1* (B). Then the co-IP and western blots were carried out as indicated. (Ci) The conserved VP motif in Cul9. (Cii) Co-IPs were performed in HEK293T cells as indicated. (Di) The immunoblots were performed in GC cells as indicated. (Dii) The immunoblots were performed in GC cells with *Yes1* knockdown or not as indicated. (E) MS/MS identified Cul9 ubiquitination residues. (F) GC cells were used for *in vivo* ubiquitination experiments. (G) The effects of *WT-Cul9* and its *K1657R* mutant in GC cells.

**Supplementary Figure 4.**
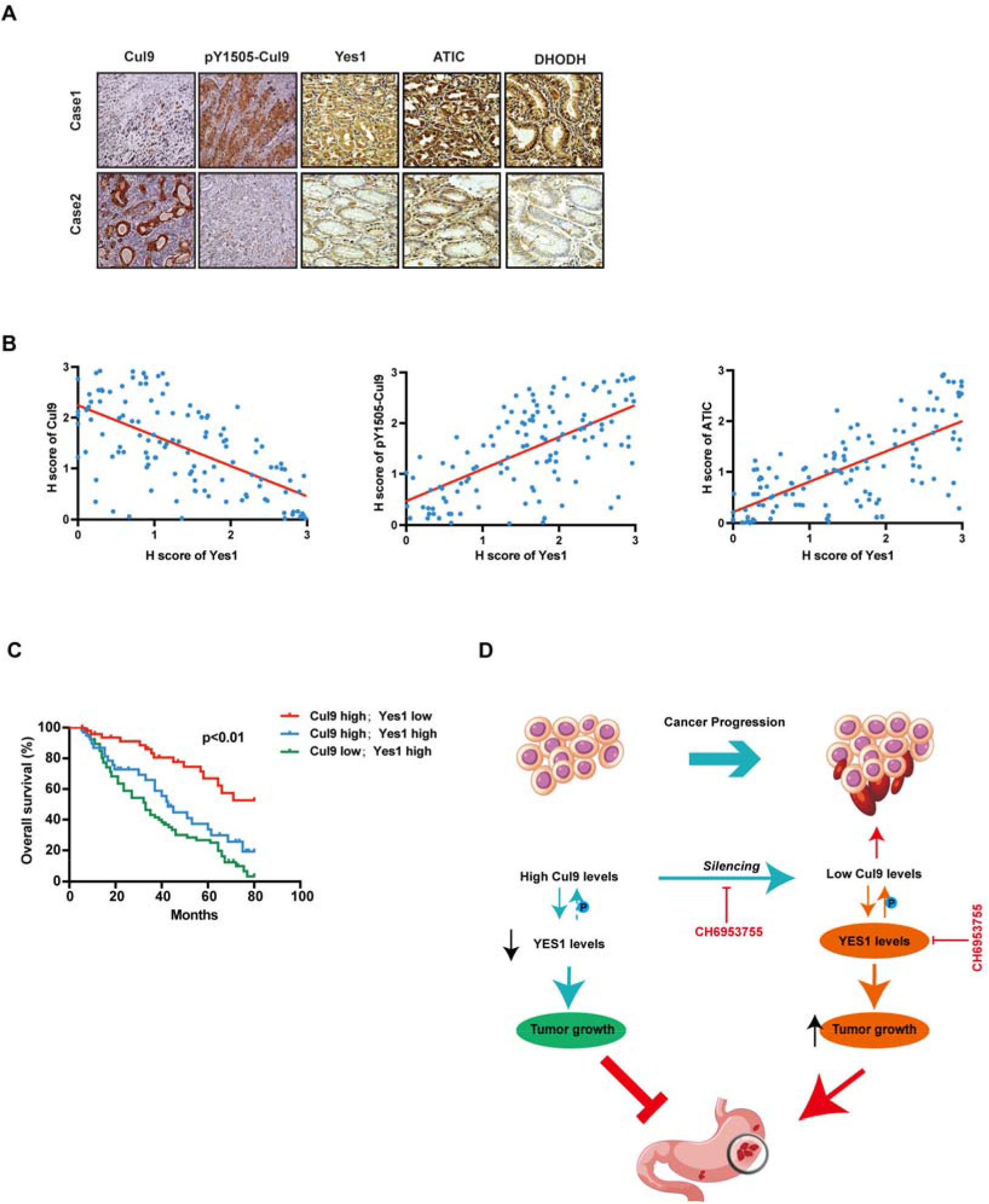
The levels of Yes1 and Cul9 are oppositely related in patients with GC. (A) IHC data. (B) The Pearson correlation assays as indicated (n =101). (C) Kaplan-Meier assays based on Cul9 and Yes1 levels in patients with GC (n=110). (D) Schematic model about the role of Yes1-Cul9 feedback axis in GC development.

**Supplementary Table S1.** Top 500 upregulated genes with high Cul9 expression in TCGA GC samples.

|  |  |  |  |  |
| --- | --- | --- | --- | --- |
| KIAA0467 | MADD | RECQL5 | MKL2 | KIAA1529 |
| C12orf51 | KIAA0913 | RPRD2 | PRRT3 | 43532 |
| USP49 | TRIM39 | H6PD | OLFML2A | KIAA0232 |
| HERC2 | CASKIN2 | TYK2 | KDM3B | SETX |
| HCFC1 | ZC3H4 | DOCK6 | SSH1 | UCKL1AS |
| GON4L | DYNC1H1 | LOC200030 | KSR1 | ?\|155060 |
| CUL7 | NR2C2 | SFRS14 | RAI1 | SPDYE6 |
| HERC1 | MTOR | BTBD9 | ZNF641 | ZMIZ1 |
| EP400 | CEP250 | GPR135 | KIAA0355 | CHD4 |
| KIAA0892 | BPTF | DOCK9 | ZFAT | MKL1 |
| WDR81 | ZXDC | NPLOC4 | OMG | ARHGEF1 |
| RPTOR | TTC21A | TRIO | GRIPAP1 | KIAA0947 |
| ABCC10 | NOTCH1 | PHRF1 | UNC45A | PAPLN |
| FBF1 | PKD1 | TBC1D13 | OTUD7B | OTUD7A |
| CRAMP1L | SLC26A1 | ZNF490 | PRICKLE4 | FAM22F |
| GOLGA3 | CLASP1 | CYTSA | FYCO1 | EZH1 |
| TAF8 | NSD1 | ZNF192 | URB1 | TULP4 |
| TUBGCP6 | DENND4B | LRIG2 | KRBA2 | TCP11L1 |
| SIN3B | MAPK8IP3 | KIAA0226 | PRIC285 | MED22 |
| EXOC7 | TNRC18 | TRRAP | SP2 | MTMR15 |
| CIC | MLL4 | C19orf44 | LOC100271836 | LOC91316 |
| MCM3AP | ZNF460 | C19orf55 | KIAA0754 | C2orf86 |
| ASB16 | ZER1 | BRD4 | ZBTB22 | WNK1 |
| PCNT | PGS1 | ZNF609 | NAV2 | C20orf165 |
| KIAA0240 | SRCAP | C2CD3 | FAM193B | ZNF366 |
| LOC283922 | PHF12 | NUP214 | KLHL36 | ZNF8 |
| TJAP1 | SYNRG | AMBRA1 | LOC619207 | SIN3A |
| MLL2 | EHMT1 | BRPF3 | CRY2 | ANKRD23 |
| KIAA1267 | EME2 | MAST4 | NOTCH4 | MEF2D |
| ZNF407 | NBPF10 | LOC100133331 | TAF1 | CCDC57 |
| LOC100170939 | ZFYVE26 | RFX1 | GIGYF1 | CSPG4 |
| ATG2A | MEGF8 | WDR88 | LLGL1 | LOC100129550 |
| ANKS1A | LZTR1 | NOL9 | TRAPPC10 | TSC1 |
| CREBBP | GTF3C1 | AKNA | DGCR8 | FRS3 |
| CHD8 | WIZ | TRIM56 | PCDHGC5 | ZNF767 |
| SETD1B | ZNF445 | ZNF687 | ALMS1 | DHX16 |
| MYO9B | STAT5B | NFATC2 | C9orf45 | CLEC16A |
| TSC2 | GFOD1 | PTPN14 | ADCY6 | LOC650623 |
| BAT2L1 | UBR2 | GNA12 | ZSCAN20 | NCOA3 |
| SYNGAP1 | SETD1A | EDC4 | KHNYN | TRIM66 |
| ZBTB37 | HSPG2 | GPATCH8 | LOC285033 | ACIN1 |
| ARHGEF11 | ZNF646 | KDM4B | CABIN1 | MYCBP2 |
| RAD54L2 | YLPM1 | SLC25A42 | RPAP1 | GMEB2 |
| CHD6 | HTT | TGFBRAP1 | CROCCL2 | ADCY4 |
| MAPKBP1 | MLL | TTBK2 | MBD5 | TENC1 |
| SRRM2 | MAML1 | BIRC6 | ASXL1 | SDHAP1 |
| NUMA1 | TRIM62 | POGZ | DCAF8 | SCAP |
| ARID1B | INPP5B | PDPR | USHBP1 | PLXNA1 |
| RAPGEF1 | PDE4DIP | C6orf106 | KIAA1530 | LENG8 |
| ZNF710 | LIMD1 | PLEKHM2 | BTBD18 | VWF |
| BTBD12 | INTS3 | USP19 | IRGQ | CNST |
| SKIV2L | LOC100129726 | CASZ1 | 43717 | MAST3 |
| MED12 | MAP3K14 | TCP11L2 | TAC4 | ACACB |
| ZZEF1 | PLCG1 | NBPF14 | KIAA1614 | LOC283314 |
| ZNF318 | GLTSCR1 | KCTD2 | STK36 | ALX3 |
| SLC35E2 | LAMA5 | FAM193A | CRTC1 | PIK3C2B |
| MICAL3 | NISCH | NDST1 | ASB1 | ZBTB20 |
| SMG6 | TLN1 | BAHD1 | DDI2 | TRIM44 |
| UBR4 | UPF1 | IGHMBP2 | TSPYL3 | TAOK2 |
| ZBTB3 | MYO18A | PGAM2 | MAP3K3 | SON |
| DIDO1 | IPO13 | ZNF496 | SNRNP200 | TBC1D16 |
| USP20 | HELZ | FAM53B | PACS1 | C20orf132 |
| LOC728989 | ZNF70 | ATXN1L | ZFHX3 | BTBD8 |
| PRR12 | SETD2 | C14orf43 | ZNF785 | CEP97 |
| C14orf49 | MACF1 | TNRC6B | NEK9 | DCTN1 |
| TMEM151B | NCOR2 | RBM33 | C14orf21 | PCNX |
| TSSK4 | SGSM2 | ZFP41 | RC3H1 | WDTC1 |
| TECPR2 | DNHD1 | CROCC | LRRK1 | BZRAP1 |
| ZFYVE20 | ZNF592 | MDN1 | EP300 | CRTC3 |
| PCNXL3 | PPM1F | GTPBP1 | BRD1 | LMTK2 |
| KIAA1310 | MYH9 | LTB4R2 | MAN2A2 | DNAH1 |
| PI4KA | SPDYE8P | SMCR8 | SYNPO | MGA |
| MLLT6 | GBF1 | GATAD2B | SH3TC1 | PMS2L11 |
| VPS39 | FAM22G | TBC1D2B | C15orf28 | USP35 |
| NEURL4 | MLL3 | INPPL1 | GDF7 | HDAC7 |
| IQSEC1 | MED13L | PI4KB | CYB561D1 | PTPN23 |
| VPS13D | C14orf4 | ITPKB | HSFX2 | ERC1 |
| TECPR1 | PHF2 | ST5 | FAM186B | DCDC2B |
| LOC389333 | SBF1 | MECP2 | POM121C | C9orf102 |
| ZBTB40 | AARS2 | UBE2O | EIF2C1 | GPR17 |
| ZNF335 | INTS1 | ZC3H7B | DCHS1 | PVRIG |
| C20orf117 | ABL1 | SHANK3 | MDC1 | PIP4K2B |
| BAT2 | ZNF500 | MTMR3 | ZNF517 | ZBTB17 |
| CELF1 | ARID1A | RUNDC2A | SEC14L1 | TRPV1 |
| TP53BP1 | TBC1D24 | ZNF236 | SIK2 | SNAPC4 |
| ZNF142 | ZHX3 | PLXNB1 | SMYD4 | TBC1D9B |
| CLCN6 | FLJ45340 | UNKL | GRLF1 | STK11IP |
| PRDM15 | PLEKHM3 | JRK | C17orf70 | SLC9A8 |
| LOC641298 | AHDC1 | HDAC4 | LRCH3 | CHD2 |
| SUPT6H | MTHFR | ELMO2 | KIAA1632 | C4orf42 |
| TLE3 | CBFA2T2 | EIF2C3 | UBE3B | ADD1 |
| SPEN | MYO9A | C22orf46 | ELL | SMG1 |
| HUWE1 | GCN1L1 | LOC100272228 | FOXK1 | MYSM1 |
| UNK | POLG | RGS12 | LOC100131193 | DIP2A |
| ASH1L | ERN1 | SCAND2 | C6orf89 | TFEB |
| C19orf71 | TEP1 | EHBP1L1 | CDK5RAP2 | PLBD2 |
| ULK1 | MLXIP | HIVEP1 | SEC16A | ZMYM3 |
| SRF | FBXL20 | KIAA0556 | RREB1 | C14orf118 |
| RBM15 | ARAP1 | FTSJD2 | MAN2C1 | LOC285740 |
| GGA3 | KIAA0195 | TSSK6 | TADA2B | SUN2 |

**Supplementary Table S2.**
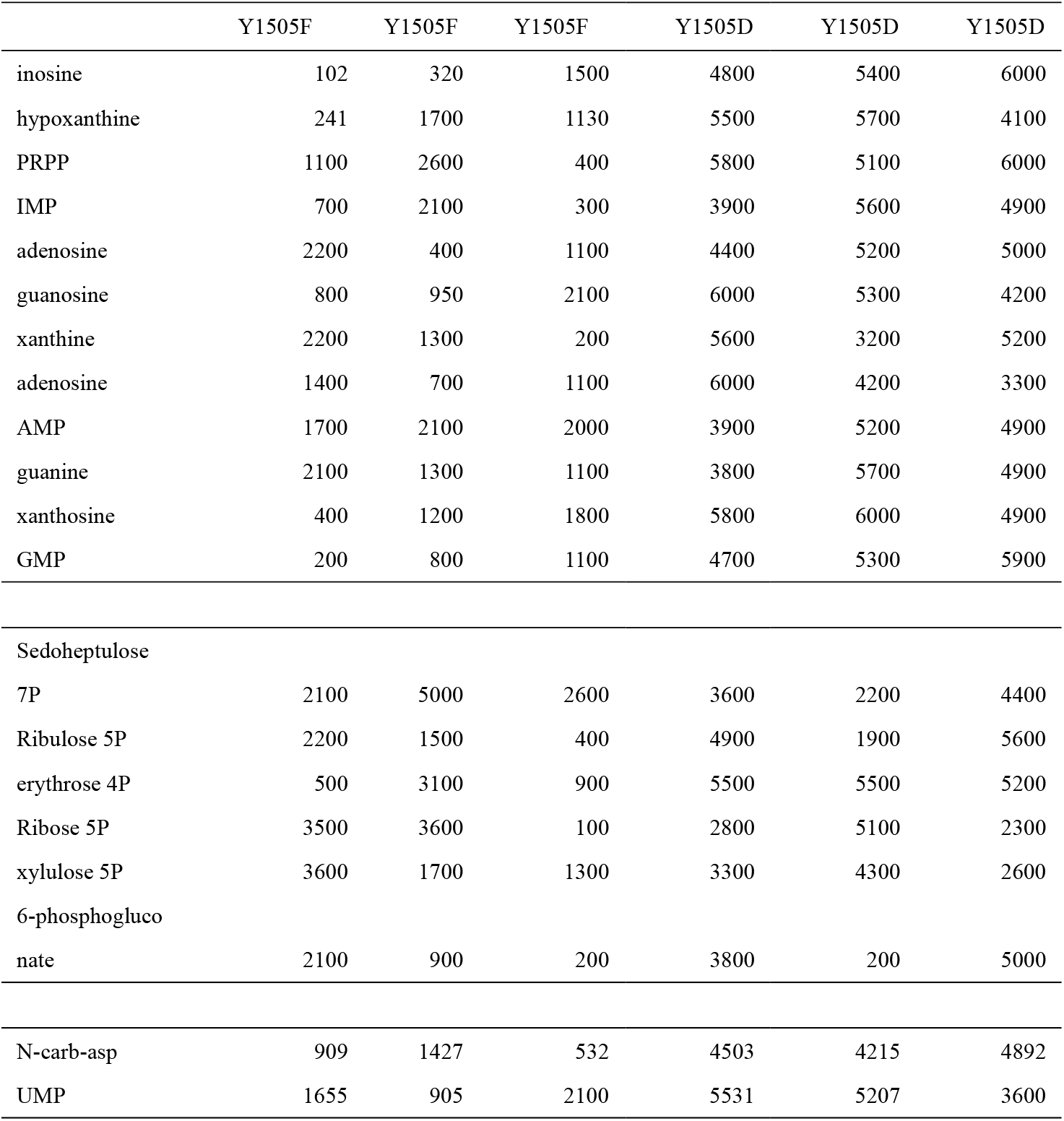
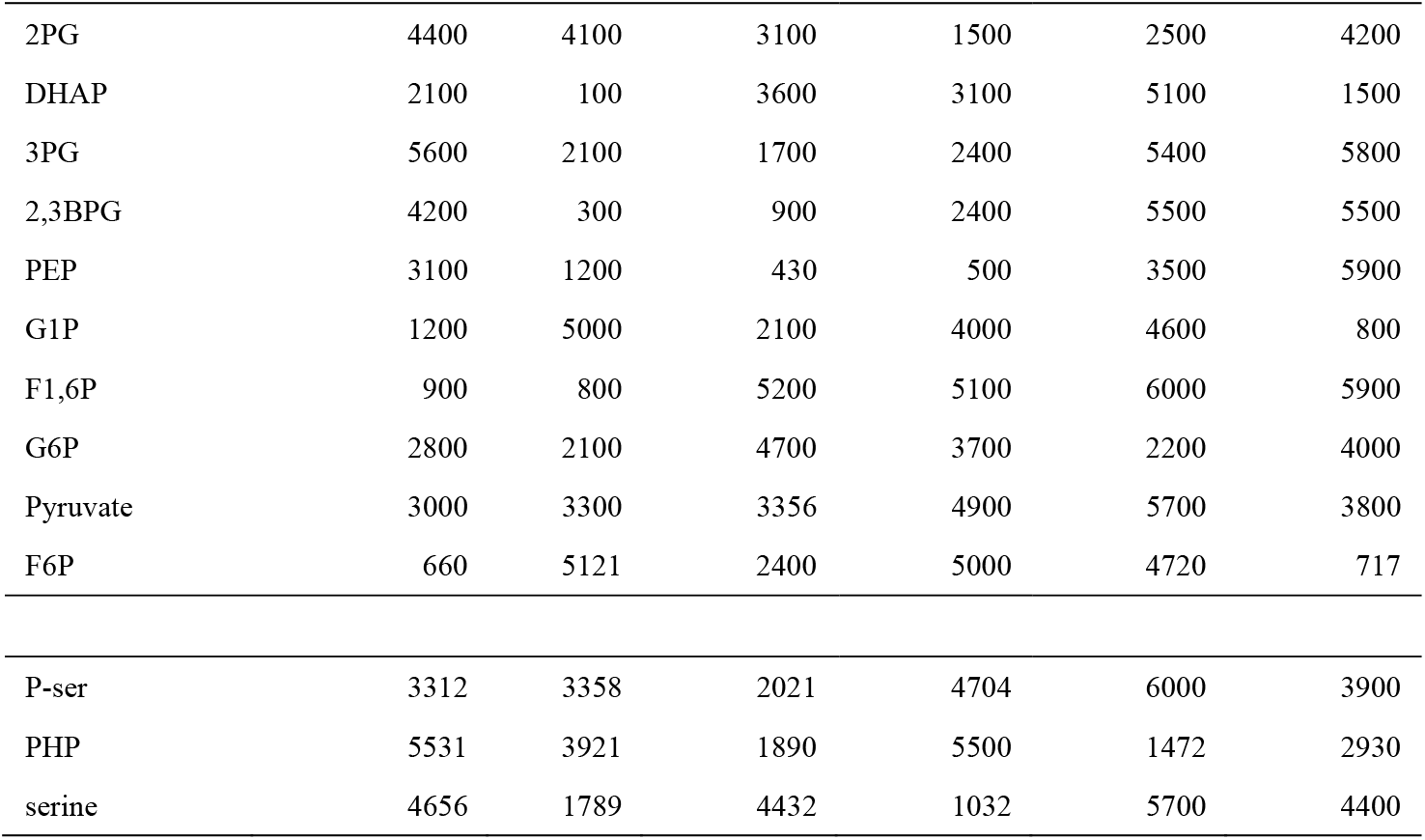
The metabolic profiling in SGC7901 cells expressing Cul9-Y1505F and Cul9-Y1505D mutant.

**Supplementary Table S3.**
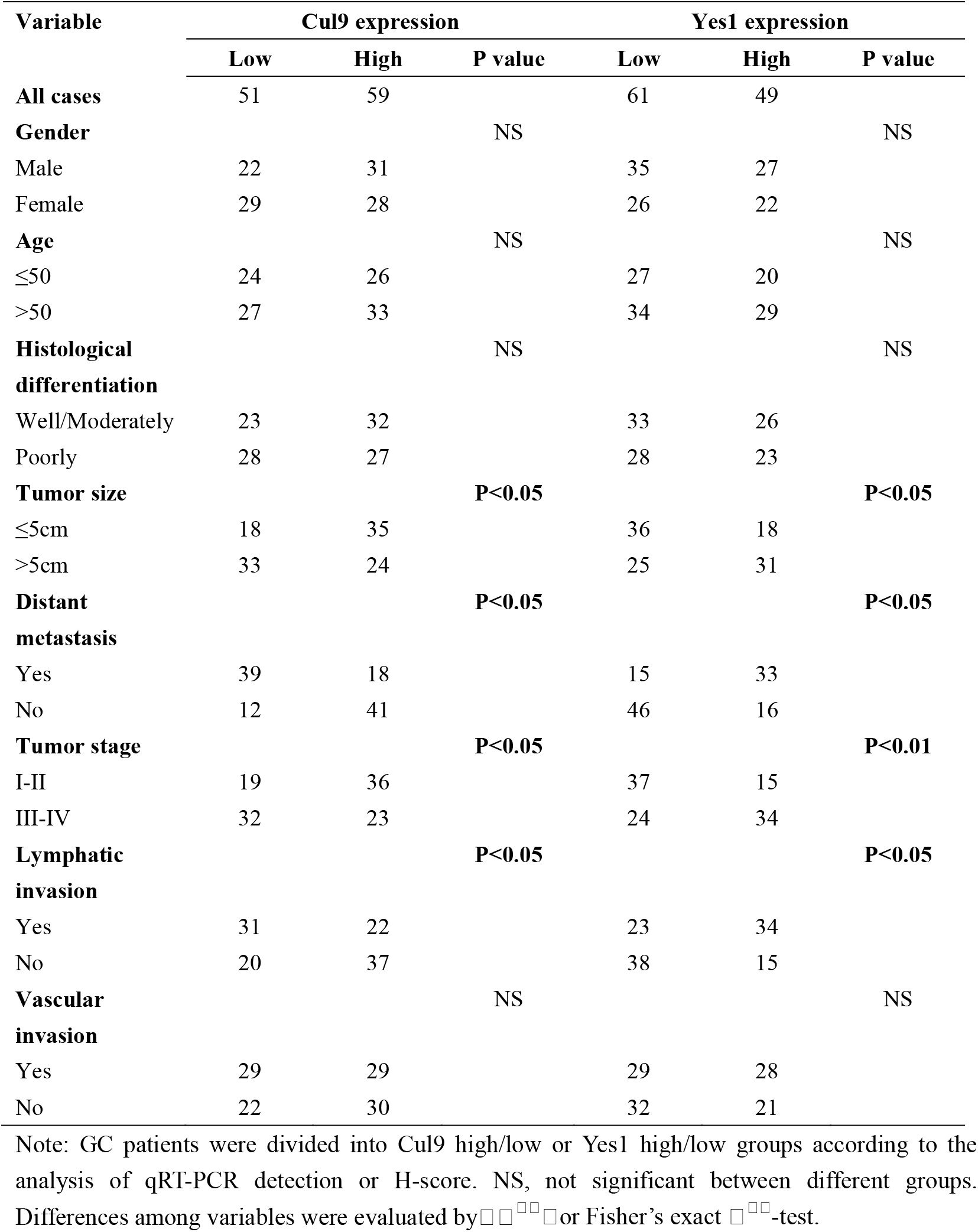
Relationship between Cul9 or Yes1 expression and clinicopathological features of GC patients (n=110)

| Variable | Cul9 expression |  |  | Yes1 expression |  |  |
| --- | --- | --- | --- | --- | --- | --- |
|  | Low | High | P value | Low | High | P value |
| **All cases** | 51 | 59 |  | 61 | 49 |  |
| **Gender** |  |  | NS |  |  | NS |
| Male | 22 | 31 |  | 35 | 27 |  |
| Female | 29 | 28 |  | 26 | 22 |  |
| **Age** |  |  | NS |  |  | NS |
| ≤50 | 24 | 26 |  | 27 | 20 |  |
| >50 | 27 | 33 |  | 34 | 29 |  |
| **Histological differentiation** |  |  | NS |  |  | NS |
| Well/Moderately | 23 | 32 |  | 33 | 26 |  |
| Poorly | 28 | 27 |  | 28 | 23 |  |
| **Tumor size** |  |  | P<0.05 |  |  | P<0.05 |
| ≤5cm | 18 | 35 |  | 36 | 18 |  |
| >5cm | 33 | 24 |  | 25 | 31 |  |
| **Distant metastasis** |  |  | P<0.05 |  |  | P<0.05 |
| Yes | 39 | 18 |  | 15 | 33 |  |
| No | 12 | 41 |  | 46 | 16 |  |
| **Tumor stage** |  |  | P<0.05 |  |  | P<0.01 |
| I-II | 19 | 36 |  | 37 | 15 |  |
| III-IV | 32 | 23 |  | 24 | 34 |  |
| **Lymphatic invasion** |  |  | P<0.05 |  |  | P<0.05 |
| Yes | 31 | 22 |  | 23 | 34 |  |
| No | 20 | 37 |  | 38 | 15 |  |
| **Vascular invasion** |  |  | NS |  |  | NS |
| Yes | 29 | 29 |  | 29 | 28 |  |
| No | 22 | 30 |  | 32 | 21 |  |

Note: GC patients were divided into Cul9 high/low or Yes1 high/low groups according to the analysis of qRT-PCR detection or H-score. NS, not significant between different groups. Differences among variables were evaluated by $\chi^2$ or Fisher's exact $\chi^2$ -test.

**Supplementary Table S4.** Univariate and multivariate analysis of factors associated with survival in GC patients (n=110)

| Variable | Univariate analysis |  | multivariate analysis |  |
| --- | --- | --- | --- | --- |
|  | HR (95% CI) | P-value | HR (95% CI) | P-value |
| **Gender** |  |  |  |  |
| Male vs. Female | 0.893(0.503-1.585) | 0.699 |  |  |
| **Age** |  |  |  |  |
| ≤50 vs. >50 | 0.776(0.434-1.389) | 0.394 |  |  |
| **Histological differentiation** |  |  |  |  |
| Well/Moderately vs. Poorly | 1.321(0.748-2.332) | 0.338 |  |  |
| **Tumor size** |  |  |  |  |
| ≤5cm vs. >5cm | 2.057(1.173-3.606) | **0.012** | 2.242(1.237-4.063) | **0.008** |
| **Distant metastasis** |  |  |  |  |
| Yes vs. No | 2.602(1.351-5.009) | **0.004** |  |  |
| **Tumor stage** |  |  |  |  |
| I-II vs. III-IV | 2.004 (1.127-3.564) | **0.018** |  |  |
| **Lymphatic invasion** |  |  |  |  |
| Yes vs. No | 1.931(1.105-3.375) | **0.021** |  |  |
| **Vascular invasion** |  |  |  |  |
| Yes vs. No | 0.814(0.444-1.494) | 0.507 |  |  |
| **Cul9 expression** |  |  |  |  |
| High vs. Low | 2.278(1.257-4.135) | **0.007** | 2.393(1.210-4.730) | **0.012** |

## References

1. Diaz S, Wang K, Sjögren B, Liu X. Roles of Cullin-RING Ubiquitin Ligases in Cardiovascular Diseases. Biomolecules 2022;12:416.

2. Li Z, Pei XH, Yan J, Yan F, Cappell KM, Whitehurst AW, Xiong Y. CUL9 mediates the functions of the 3M complex and ubiquitylates survivin to maintain genome integrity. Mol Cell 2014;54:805–19.

3. Fouad S, Wells OS, Hill MA, D’Angiolella V. Cullin Ring Ubiquitin Ligases (CRLs) in Cancer: Responses to Ionizing Radiation (IR) Treatment. Front Physiol 2019;10:1144.

4. Seipel K, Marques MT, Bozzini MA, Meinken C, Mueller BU, Pabst T. Inactivation of the p53-KLF4-CEBPA Axis in Acute Myeloid Leukemia. Clin Cancer Res 2016;22:746–56.

5. Skaar JR, Arai T, DeCaprio JA. Dimerization of CUL7 and PARC is not required for all CUL7 functions and mouse development. Mol Cell Biol. 2005 Jul;25(13):5579–89.

6. Pei XH, Bai F, Li Z, Smith MD, Whitewolf G, Jin R, Xiong Y. Cytoplasmic CUL9/PARC ubiquitin ligase is a tumor suppressor and promotes p53-dependent apoptosis. Cancer Res 2011;71:2969–77.

7. Li Z, Xiong Y. Cytoplasmic E3 ubiquitin ligase CUL9 controls cell proliferation, senescence, apoptosis and genome integrity through p53. Oncogene 2017;36:5212–5218.

8. Indovina P, Forte IM, Pentimalli F, Giordano A. Targeting SRC Family Kinases in Mesothelioma: Time to Upgrade. Cancers (Basel) 2020;12:1866.

9. Amata I, Maffei M, Pons M. Phosphorylation of unique domains of Src family kinases. Front Genet 2014;5:181.

10. Arbesú M, Maffei M, Cordeiro TN, Teixeira JM, Pérez Y, Bernadó P, Roche S, Pons M. The Unique Domain Forms a Fuzzy Intramolecular Complex in Src Family Kinases. Structure 2017;25:630–640.e4.

11. Kyo S, Sada K, Qu X, Maeno K, Miah SM, Kawauchi-Kamata K, Yamamura H. Negative regulation of Lyn protein-tyrosine kinase by c-Cbl ubiquitin-protein ligase in Fc epsilon RI-mediated mast cell activation. Genes Cells 2003;8:825–36.

12. Chung IC, Yuan SN, OuYang CN, Lin HC, Huang KY, Chen YJ, Chung AK, Chu CL, Ojcius DM, Chang YS, Chen LC. Src-family kinase-Cbl axis negatively regulates NLRP3 inflammasome activation. Cell Death Dis 2018;9:1109.

13. Fang Z, Yin S, Sun R, Zhang S, Fu M, Wu Y, Zhang T, Khaliq J, Li Y. miR-140-5p suppresses the proliferation, migration and invasion of gastric cancer by regulating YES1. Mol Cancer 2017;16:139.

14. Hong X, Huang H, Qiu X, Ding Z, Feng X, Zhu Y, Zhuo H, Hou J, Zhao J, Cai W, Sha R, Hong X, Li Y, Song H, Zhang Z. Targeting posttranslational modifications of RIOK1 inhibits the progression of colorectal and gastric cancers. Elife 2018;7:e29511.

15. Kim JS, Shin IS, Shin NR, Nam JY, Kim C. Genome-wide analysis of DNA methylation and gene expression changes in an ovalbumin-induced asthma mouse model. Mol Med Rep 2020;22:1709–1716.

16. Wu Y, Yun D, Zhao Y, Wang Y, Sun R, Yan Q, Zhang S, Lu M, Zhang Z, Lu D, Li Y. Down regulation of RNA binding motif, single-stranded interacting protein 3, along with up regulation of nuclear HIF1A correlates with poor prognosis in patients with gastric cancer. Oncotarget 2017;8:1262–1277.

17. Ohuchida K, Mizumoto K, Ishikawa N, Fujii K, Konomi H, Nagai E, Yamaguchi K, Tsuneyoshi M, Tanaka M. The role of S100A6 in pancreatic cancer development and its clinical implication as a diagnostic marker and therapeutic target. Clin Cancer Res 2005;11:7785–93.

18. Hu Y, Chen H, Zhang L, Lin X, Li X, Zhuang H, Fan H, Meng T, He Z, Huang H, Gong Q, Zhu D, Xu Y, He P, Li L, Feng D. The AMPK-MFN2 axis regulates MAM dynamics and autophagy induced by energy stresses. Autophagy. 2021;17:1142–1156.

19. Han SH, Korm S, Han YG, Choi SY, Kim SH, Chung HJ, Park K, Kim JY, Myung K, Lee JY, Kim H, Kim DW. GCA links TRAF6-ULK1-dependent autophagy activation in resistant chronic myeloid leukemia. Autophagy 2019;15:2076–2090.

20. Zhu S, Soutto M, Chen Z, Peng D, Romero-Gallo J, Krishna US, Belkhiri A, Washington MK, Peek R, El-Rifai W. *Helicobacter pylori-*induced cell death is counteracted by NF-κB-mediated transcription of DARPP-32. Gut 2017;66:761–762.

21. Nagy TA, Frey MR, Yan F, Israel DA, Polk DB, Peek RM Jr. Helicobacter pylori regulates cellular migration and apoptosis by activation of phosphatidylinositol 3-kinase signaling. J Infect Dis 2009;199:641–51.

22. Franco AT, Johnston E, Krishna U, Yamaoka Y, Israel DA, Nagy TA, Wroblewski LE, Piazuelo MB, Correa P, Peek RM Jr. Regulation of gastric carcinogenesis by Helicobacter pylori virulence factors. Cancer Res 2008;68:379–87.

23. Ma F, Zhu Y, Liu X, Zhou Q, Hong X, Qu C, Feng X, Zhang Y, Ding Q, Zhao J, Hou J, Zhong M, Zhuo H, Zhong L, Ye Z, Xie W, Liu Y, Xiong Y, Chen H, Piao D, Sun B, Gao Z, Li Q, Zhang Z, Qiu X, Zhang Z. Dual-Specificity Tyrosine Phosphorylation-Regulated Kinase 3 Loss Activates Purine Metabolism and Promotes Hepatocellular Carcinoma Progression. Hepatology 2019;70:1785–1803.

24. Ben-Sahra I, Hoxhaj G, Ricoult SJH, Asara JM, Manning BD. mTORC1 induces purine synthesis through control of the mitochondrial tetrahydrofolate cycle. Science 2016;351:728–733.

25. Shen Y, Zhang H, Yao S, Su F, Wang H, Yin J, Fang Y, Tan L, Zhang K, Fan X, Zhong M, Zhou Q, He J, Zhang Z. Methionine oxidation of CLK4 promotes the metabolic switch and redox homeostasis in esophageal carcinoma via inhibiting MITF selective autophagy. Clin Transl Med 2022;12:e719.

26. Du Q, Luu PL, Stirzaker C, Clark SJ. Methyl-CpG-binding domain proteins: readers of the epigenome. Epigenomics 2015;7:1051–73.

27. Feng X, Zhang H, Meng L, Song H, Zhou Q, Qu C, Zhao P, Li Q, Zou C, Liu X, Zhang Z. Hypoxia-induced acetylation of PAK1 enhances autophagy and promotes brain tumorigenesis via phosphorylating ATG5. Autophagy 2021;17:723–742.

28. Song H, Feng X, Zhang M, Jin X, Xu X, Wang L, Ding X, Luo Y, Lin F, Wu Q, Liang G, Yu T, Liu Q, Zhang Z. Crosstalk between lysine methylation and phosphorylation of ATG16L1 dictates the apoptosis of hypoxia/reoxygenation-induced cardiomyocytes. Autophagy 2018;14:825–844.

29. Feng X, Jia Y, Zhang Y, Ma F, Zhu Y, Hong X, Zhou Q, He R, Zhang H, Jin J, Piao D, Huang H, Li Q, Qiu X, Zhang Z. Ubiquitination of UVRAG by SMURF1 promotes autophagosome maturation and inhibits hepatocellular carcinoma growth. Autophagy 2019;15:1130–1149.

30. Vainshtein A, Grumati P. Selective Autophagy by Close Encounters of the Ubiquitin Kind. Cells 2020;9:2349.

31. Lau K, Podolec R, Chappuis R, Ulm R, Hothorn M. Plant photoreceptors and their signaling components compete for COP1 binding via VP peptide motifs. EMBO J 2019;38:e102140.

32. Creeden JF, Alganem K, Imami AS, Brunicardi FC, Liu SH, Shukla R, Tomar T, Naji F, McCullumsmith RE. Kinome Array Profiling of Patient-Derived Pancreatic Ductal Adenocarcinoma Identifies Differentially Active Protein Tyrosine Kinases. Int J Mol Sci 2020;21:8679.

33. Feng X, Ma D, Zhao J, Song Y, Zhu Y, Zhou Q, Ma F, Liu X, Zhong M, Liu Y, Xiong Y, Qiu X, Zhang Z, Zhang H, Zhao Y, Zhang K, Hong X, Zhang Z. UHMK1 promotes gastric cancer progression through reprogramming nucleotide metabolism. EMBO J. 2020;39:e102541.

34. Zhou Q, Lin M, Feng X, Ma F, Zhu Y, Liu X, Qu C, Sui H, Sun B, Zhu A, Zhang H, Huang H, Gao Z, Zhao Y, Sun J, Bai Y, Jin J, Hong X, Zou C, Zhang Z. Targeting CLK3 inhibits the progression of cholangiocarcinoma by reprogramming nucleotide metabolism. J Exp Med. 2020;217:e20191779.

35. Ali ES, Sahu U, Villa E, O’Hara BP, Gao P, Beaudet C, Wood AW, Asara JM, Ben-Sahra I. ERK2 Phosphorylates PFAS to Mediate Posttranslational Control of De Novo Purine Synthesis. Mol Cell 2020;78:1178–1191.e6.

36. Ben-Sahra I, Howell JJ, Asara JM, Manning BD. Stimulation of de novo pyrimidine synthesis by growth signaling through mTOR and S6K1. Science 2013;339:1323–8.

37. Merhi A, Delrée P, Marini AM. The metabolic waste ammonium regulates mTORC2 and mTORC1 signaling. Sci Rep 2017;7:44602.

38. Hartmann T, Xu X, Kronast M, Muehlich S, Meyer K, Zimmermann W, Hurwitz J, Pan ZQ, Engelhardt S, Sarikas A. Inhibition of Cullin-RING E3 ubiquitin ligase 7 by simian virus 40 large T antigen. Proc Natl Acad Sci U S A 2014;111:3371–6.

39. Srivastava S, Sahu U, Zhou Y, Hogan AK, Sathyan KM, Bodner J, Huang J, Wong KA, Khalatyan N, Savas JN, Ntziachristos P, Ben-Sahra I, Foltz DR. NOTCH1-driven UBR7 stimulates nucleotide biosynthesis to promote T cell acute lymphoblastic leukemia. Sci Adv 2021;7:eabc9781.

40. Mohapatra B, Ahmad G, Nadeau S, Zutshi N, An W, Scheffe S, Dong L, Feng D, Goetz B, Arya P, Bailey TA, Palermo N, Borgstahl GE, Natarajan A, Raja SM, Naramura M, Band V, Band H. Protein tyrosine kinase regulation by ubiquitination: critical roles of Cbl-family ubiquitin ligases. Biochim Biophys Acta 2013;1833:122–39.

41. Rao N, Dodge I, Band H. The Cbl family of ubiquitin ligases: critical negative regulators of tyrosine kinase signaling in the immune system. J Leukoc Biol 2002;71:753–63.

42. Bao J, Gur G, Yarden Y. Src promotes destruction of c-Cbl: implications for oncogenic synergy between Src and growth factor receptors. Proc Natl Acad Sci U S A. 2003;100:2438–43.

43. Fan S, Weight CM, Luissint AC, Hilgarth RS, Brazil JC, Ettel M, Nusrat A, Parkos CA. Role of JAM-A tyrosine phosphorylation in epithelial barrier dysfunction during intestinal inflammation. Mol Biol Cell 2019;30:566–578.

44. Mcheik S, Aptecar L, Coopman P, D’Hondt V, Freiss G. Dual Role of the PTPN13 Tyrosine Phosphatase in Cancer. Biomolecules. 2020;10:1659.

45. Hamyeh M, Bernex F, Larive RM, Naldi A, Urbach S, Simony-Lafontaine J, Puech C, Bakhache W, Solassol J, Coopman PJ, Hendriks WJAJ, Freiss G. PTPN13 induces cell junction stabilization and inhibits mammary tumor invasiveness. Theranostics. 2020;10:1016–1032.

46. Glondu-Lassis M, Dromard M, Lacroix-Triki M, Nirdé P, Puech C, Knani D, Chalbos D, Freiss G. PTPL1/PTPN13 regulates breast cancer cell aggressiveness through direct inactivation of Src kinase. Cancer Res 2010;70:5116–26.

47. Kazanets A, Shorstova T, Hilmi K, Marques M, Witcher M. Epigenetic silencing of tumor suppressor genes: Paradigms, puzzles, and potential. Biochim Biophys Acta 2016;1865:275–88.

48. Wood KH, Zhou Z. Emerging Molecular and Biological Functions of MBD2, a Reader of DNA Methylation. Front Genet 2016;7:93.

49. de Pins B, Mendes T, Giralt A, Girault JA. The Non-receptor Tyrosine Kinase Pyk2 in Brain Function and Neurological and Psychiatric Diseases. Front Synaptic Neurosci 2021;13:749001.

